# Embryonic lymphocytes contribute to a genetic form of autoimmune inflammation

**DOI:** 10.1101/2023.11.24.568475

**Authors:** Sara Cascione, Elena Fontana, Rebecca Scarfò, Rosita Rigoni, Valentina Capo, Elena Draghici, Kerry Dobbs, Luigi D. Notarangelo, Anna Villa, Andrea Ditadi

**Author notes:** Corresponding author Andrea Ditadi San Raffaele Telethon Institute for Gene Therapy IRCCS San Raffaele Scientific Institute Via Olgettina, 58 20132 Milano (Italy).

## Abstract

Omenn Syndrome (OS) is a rare hematological disorder, caused by hypomorphic mutations in genes involved in B-/T-cell receptor (BCR/TCR) rearrangement that result in impaired lymphocyte development and immunodeficiency. Notwithstanding, few T-cell clones enriched in self-reactive specificities expand in peripheral tissues, where they trigger severe inflammation and autoimmune reactions. Interestingly, residual OS lymphocytes display characteristics proper of embryonic lymphocytes that emerge before, and independently from, hematopoietic stem cells (HSCs). This prompted us to hypothesize whether OS autoreactive T-cells are generated in the embryo independently from HSCs. Here we show that in the *Rag2^R229Q/R229Q^* OS mouse model, embryonic but not adult bone marrow-derived hematopoietic progenitors can generate T-cells. T-lymphopoiesis can be rescued in adult OS blood progenitors via their Lin28-mediated reprogramming to an embryonic-like state. Remarkably, when transplanted in immunodeficient mice, embryonic-like OS progenitors trigger tissue morphological alterations and inflammation in the large intestine of the recipients, recapitulating the typical OS inflammatory phenotype. Our study describes the previously unappreciated contribution of embryonic progenitors to the pool of autoreactive infilitrating T-cells, providing a novel platform for both the detailed study of human autoimmune disorders and the design of more targeted therapies.

## Introduction

Severe combined immunodeficiencies (SCID) define a heterogeneous group of genetic hematological diseases that manifest in the first years of life with failure to thrive and recurrent infections, due to an impaired development of the adaptive immune system^1^. Omenn Syndrome (OS) is characterized by an unconventional combination of immunodeficiency and autoimmune-ike manifestations^2^. In fact, OS is associated with multiple genetic abnormalities that severely reduce, but not completely impair, lymphocyte development. Most of OS cases are caused by hypomorphic mutations in Recombination Activating Genes (RAG) 1 and 2, thus affecting V(D)J recombination, while null mutations in the same genes lead to RAG-SCID^1^. A minority of OS forms are caused by mutations affecting other players of the V(D)J recombination machinery as well as signaling transduction receptors^3–5^. Of note, OS patients carrying *IL7R* mutations have also been described^6^. Allogenic bone marrow transplantation represents the only available cure for OS patients, but this treatment can be ineffective, as residual activated lymphocytes may reject the transplanted cells^7^. Characterizing the residual autoreactive lymphocytes is of primary importance for the prognosis of OS and other leaky SCID patients. Most OS patients lack circulating B-cells and are not able to mount an effective humoral response to foreign antigens; nevertheless, high serum levels of self-reactive immunoglobulins are often detected and correlate with severe autoimmune manifestations^4^. The presence of T-cells is mostly confined to peripheral locations (e.g. skin and intestine)^8^, hinting at a non-random tissue migration of these cells. Interestingly, OS tissue-infiltrating T-cells display a highly restricted and shared T-cell receptor (TCR) usage across patients, which suggests a conserved and recurrent program of antigen receptor rearrangement^9^. OS T-cells in peripheral tissues express markers of activation (such as CD45RO and HLA-DR), but display a depressed response to foreign antigens, thus not contributing to a functional immune response^10^. In contrast, the TCR clonotypes shared among OS patients are enriched for self-reactive specificities, thus making the residual OS T-cells prone to get activated against the tissues they populate^11–13^. Collectively, these and other characteristics such as oligoclonality, IL7-independence and cytokine production, are very similar to those of embryonic lymphocytes, that emerge before, and independently from, hematopoietic stem cells (HSCs)^14–18^. In fact, recent studies have characterized the existence and the clinical relevance of immune cells that are restrictively generated during prenatal life^19,20^. Among the best characterized embryonic lymphocytes, B1 and marginal zone (MZ) Bcells^21^, predominantly present in murine peritoneal cavities and spleen, are known to have an embryonic-restricted origin, with post-natal HSCs only minimally contributing to such lineages^22,23^. Similarly, genetic mouse models have revealed that IL-17-producing γδ T-cells, as Vγ3 Dendritic Epidermal T cells (DETC) or lung Vγ6 cells, originate exclusively from embryonic hematopoietic progenitors and clonally self-renew in the adult^24–26^. The inability of adult HSCs to contribute to early embryonic lineages is likely due to a specific gene expression landscape that locks peculiar cellular development to a specific developmental window. Indeed, embryonic-derived cells are generated via a Lin28^+^ progenitor intermediate. The transcription factor Lin28 is expressed in erythroid, megakaryocytic and lymphoid progenitors during embryonic development, but not in the adult counterpart^27–29^. Remarkably, Lin28 ectopic expression reprograms hematopoietic stem/progenitor cells (HSPCs) from adult bone marrow to an embryonic-like state, enabling their differentiation into embryonic-specific lineages^29^. In this work, we demonstrated the contribution of embryonic lymphocytes to OS autoimmune-like manifestations. Using the *Rag2^R229Q/R229Q^* mouse model that faithfully recapitulates OS^30^, we showed that embryonic-like T-cells can recapitulate OS inflammation in peripheral tissues when transplanted in immunodeficient mice. Similarly to what observed in OS mice, human embryonic-like progenitors generated from OS patient-derived induced pluripotent stem cells (iPSC) also give rise to T-cells *in vitro*. These studies provide a new paradigm for the study of autoimmune disorders, as well as a novel platform for designing more targeted therapies.

## Results

### Residual *Rag2^R229Q/R229Q^* B lymphocytes are mostly B1 and MZ B-cells

In the *Rag2^R229Q/R229Q^* model (herein indicated as *Rag2^R229Q^*), B-cell progenitors in the bone marrow (BM) display a developmental arrest at the early pro-B stage, and mature B-cells are absent in the circulation^30^. Nevertheless, few IgM^+^ B-cells are present in tissues and display a high expression of activation markers^30,31^. To further characterize residual B-lymphocytes in *Rag2^R229Q^* mice, we analyzed the spleen and the peritoneal cavity (PerC), two of the most important peripheral B-cell reservoirs^32^. In adult *Rag2^R229Q^*mice, the frequency of total CD19^+^IgM^+^ B-cells amongst hematopoietic cells in both spleen and PerC was reduced by 27- and 3-folds respectively, compared to *Rag2^+/+^* mice (Figures S1A and S1B). We confirmed that in the adult *Rag2^R229Q^* spleen, CD21^low^CD23^+^ follicular (FO) B-cells were extremely rare (16-fold reduction with respect to *Rag2^+/+^*), while we detected a residual sizable population of marginal zone (MZ) CD21^+^CD23^neg^ B-cells (Figures 1A and 1B), as previously described^30^. In addition, in the adult *Rag2^R229^*^Q^ PerC, the CD19^mid^B220^high^ conventional B-2 B-cell fraction was severely depleted (17-fold reduction), whereas CD19^high^B220^neg/mid^ innate B-1 B-cell proportion was increased by 1.7-fold compared to *Rag2^+/+^* mice (Figures 1C and 1D). The residual presence of specific subtypes of B-cells might suggest that selected programs of lymphopoiesis can develop in OS mice. Since MZ and B1 B-cells are known to be predominantly, if not entirely, generated from early embryonic progenitors^21,23^, their residual and selective presence suggests that the maturation of pre-natal B-cell progenitors is compatible with OS *Rag2^R229Q^* hypomorphic mutation.

**Figure 1.**
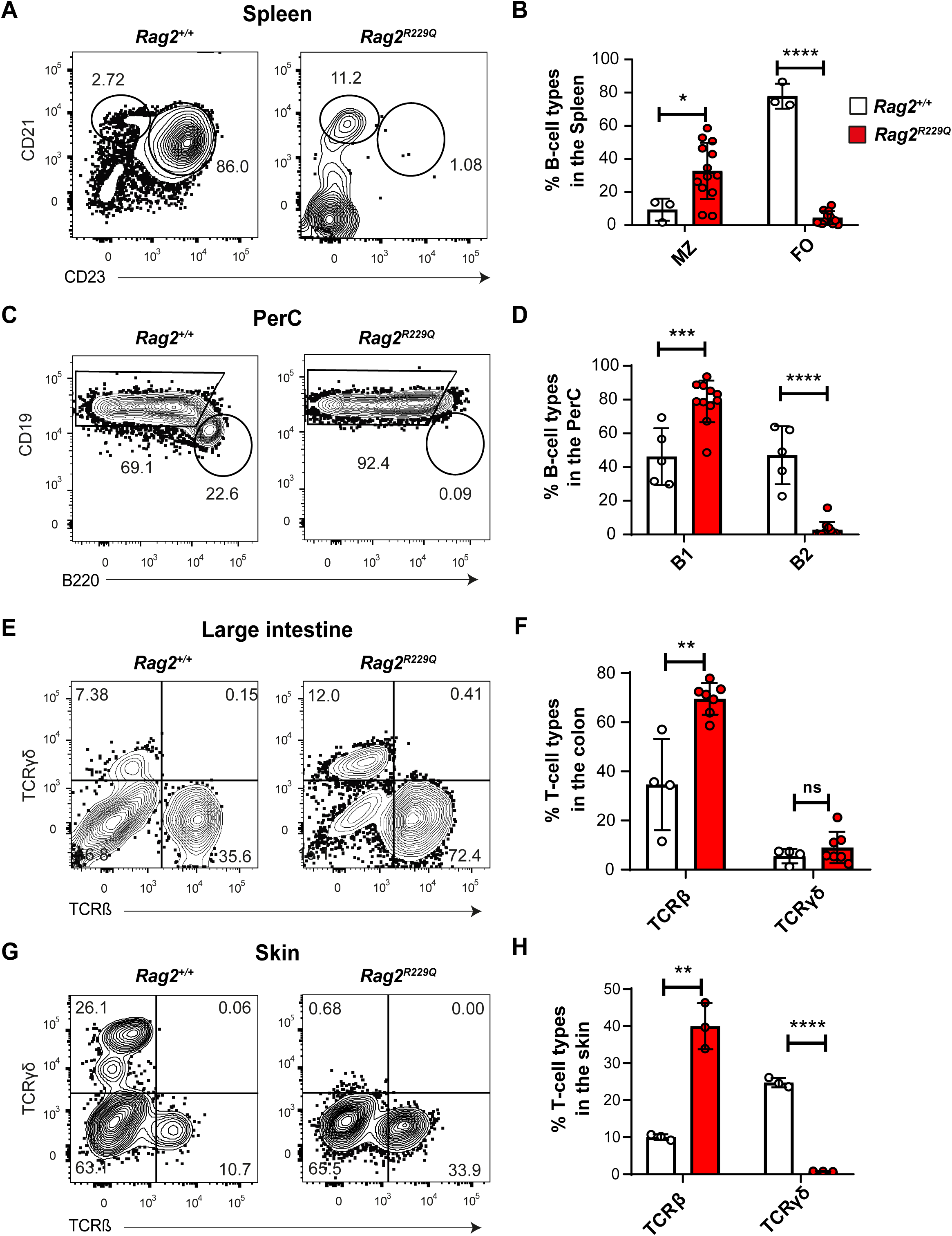
Innate B-cells and TCRβ T-cells are present in the peripheral tissues of *Rag2^R229Q^* Omenn Syndrome mice. (A) Representative flow cytometric analysis of CD21^+^CD23^neg^ MZ and CD21^+^CD23^+^ FO sub-populations of CD19^+^IgM^+^ B-cells in the spleen of adult *Rag2^R229Q^* and *Rag2^+/+^*mice, gated on FSC/SSC/7AAD^neg^/CD45^+^ cells (gating strategy shown in Figure S5A). (B) Proportion of CD21^+^CD23^neg^ MZ and CD21^+^CD23^+^ FO B-cells in the spleen of *Rag2^+/+^* (white) and *Rag2^R229Q^* (red) mice as analyzed by FACS in (A). n=3 *Rag2^+/+^* and n=13 *Rag2^R229Q^* mice from 3 independent litters. (C) Representative flow cytometric analysis of CD19^high^B220^neg/mid^ innate embryonic B1 and CD19^mid^B220^high^ conventional B2 cells sub-populations within the CD19^+^IgM^+^ B-cell of the peritoneal cavity (PerC) of *Rag2^R229Q^* and *Rag2^+/+^*mice. Cells are gated on FSC/SSC/7AAD^neg^/CD45^+^ cells (gating strategy shown in Figure S5B). (D) Frequency of innate B1 and conventional B2 in the PerC of *Rag2^+/+^* (white) and *Rag2^R229Q^* (red) mice as analyzed by FACS in (C). n=5 *Rag2^+/+^* and n=11 *Rag2^R229Q^* mice from 4 independent litters. (E) Representative flow cytometric analysis of TCRβ and TCRγδ T-cells in the large intestine of *Rag2^R229Q^*and *Rag2^+/+^* mice, gated on FSC/SSC/CD45^+^/7AAD^neg^ (gating strategy shown in Figure S5C). Proportions are shown in panel (F). n=4 *Rag2^+/+^*and n=7 *Rag2^R229Q^* mice from 2 independent litters. (G) Representative flow cytometric analysis of β and γδ T-cells in the skin of *Rag2^R229Q^*and *Rag2^+/+^* mice, gated on FSC/SSC/CD45^+^/7AAD^neg^. Proportions are shown in (H). n=3 *Rag2^+/+^* and n=3 *Rag2^R229Q^* mice from 2 independent litters. p-values are for unpaired Student’s t-test. *p<0.05, **p<0.01, ***p<0.001, **** p<0.0001.

### T-lymphocytes are expanded in the peripheral tissues of *Rag2^R229Q^* mice

As the major clinical OS manifestations are caused by an aberrant activation of T-cells in the periphery, with preferential localization in skin and gut^8,12,33,34^, we looked more closely at peripheral T-cell compartments in adult *Rag2^R229Q^* OS mice. In *Rag2^R229Q^* large intestines, the overall proportion of T-lymphocytes amongst hematopoietic cells was increased by 2-fold compared to *Rag2^+/+^* controls (Figure S1C) and composed by both TCRβ T-cells, representing the major T-cell type with a 2-fold increased frequency compared to *Rag2^+/+^*, and TCRγδ T-cells (Figures 1E and 1F). In contrast, the skin of *Rag2^R229^*^Q^ mice did not display any significant difference in T-cells abundance (Figure S1D). However, we confirmed the predominant presence of the TCRβ over the TCRγδ T-cells (Figures 1G and 1H), as we previously reported^30^. Remarkably, despite the presence of TCRβ T-cells in the periphery, only few, if any, CD4^+^CD8^+^ double positive (DP) thymocytes were observed in the *Rag2^R229^*^Q^ thymus (Figures S1E and S1F), in line with previous observations^30^. The residual presence of T-cells in both skin and large intestine in the absence of DP thymic T-lymphoid progenitors suggests an alternative and/or complementary origin of the residual T-cells in OS mice.

### Embryonic but not adult *Rag2^R229^*^Q^ hematopoietic progenitors give rise to DP and CD3^+^ T-cells *ex vivo*

Given these observations about peripheral lymphocytes and considering the similarity between residual OS and embryonic-derived lymphocytes, in terms of peripheral location, short CDR3 and self-reactivity^30,35^, we decided to investigate the origin of the residual T-cells present in the *Rag2^R229Q^* mouse model. To this end, we tested the T-cell potential of *Rag2^R229Q^* adult and early embryonic hematopoietic progenitors *ex vivo,* using a standard T-cell differentiation assay based on OP9-DLL1 co-culture^36^ (Figure S2A). Under these conditions, lineage negative (Lin^neg^) HSPCs isolated from the BM of 12-14 weeks old Rag2^+/+^ mice gave rise to all the CD4^neg^CD8^neg^ double negative (DN) populations (Figure S2B, top) and successfully generated DP progenitors (Figures 2A and 2B) and more mature cells expressing the TCR co-receptor CD3 (Figure S2C, top). In contrast, *Rag2^R229Q^*lymphocytes could not progress beyond the CD44^neg^CD25^+^ DN3 stage (Figure S2B) and failed to generate DP and CD3^+^ cells in culture (Figures 2A, 2B and S2C), thus recapitulating the developmental block of *Rag2^R229Q^* thymocytes at DN3 stage observed *in vivo*^30^. This severe impairment of *Rag2^R229Q^* adult progenitors in generating DP T-cells supports the hypothesis of an alternative or complementary source to BM HSPCs for the OS T-cells. We next interrogated the impact of the *Rag2^R229Q^*mutation on embryonic lymphopoiesis. Since the first hematopoietic progenitors with lymphoid potential are found in the yolk sac (YS) and Para-aortic splanchnopleure (P-Sp) before the emergence of the first HSCs^37^, we tested the ability of *Rag2^R229Q^* and *Rag2^+/+^* E9.5 YS- and P-Sp-derived cells to differentiate into T-cells. In stark contrast to adult HSPCs, *Rag2^R229Q^* YS- and P-Sp-derived cells transitioned from the DN3 to the DN4 stage (Figure S2B) and gave rise to DP T-cells in similar proportions to *Rag2^+/+^* embryos’ cultures (Figure 2C and 2D). Moreover, in contrast to the BM HSPCs culture, *Rag2^R229Q^*E9.5 cells could generate CD3^+^ T-cells (Figure S2C), albeit at a low frequency (Figure S2D). Taken together, these results strongly suggest that the *Rag2^R229Q^* OS mutation does not impair the progression of early embryonic lymphoid progenitors to the DP stage and the generation of mature CD3^+^ T-cells.

**Figure 2.**
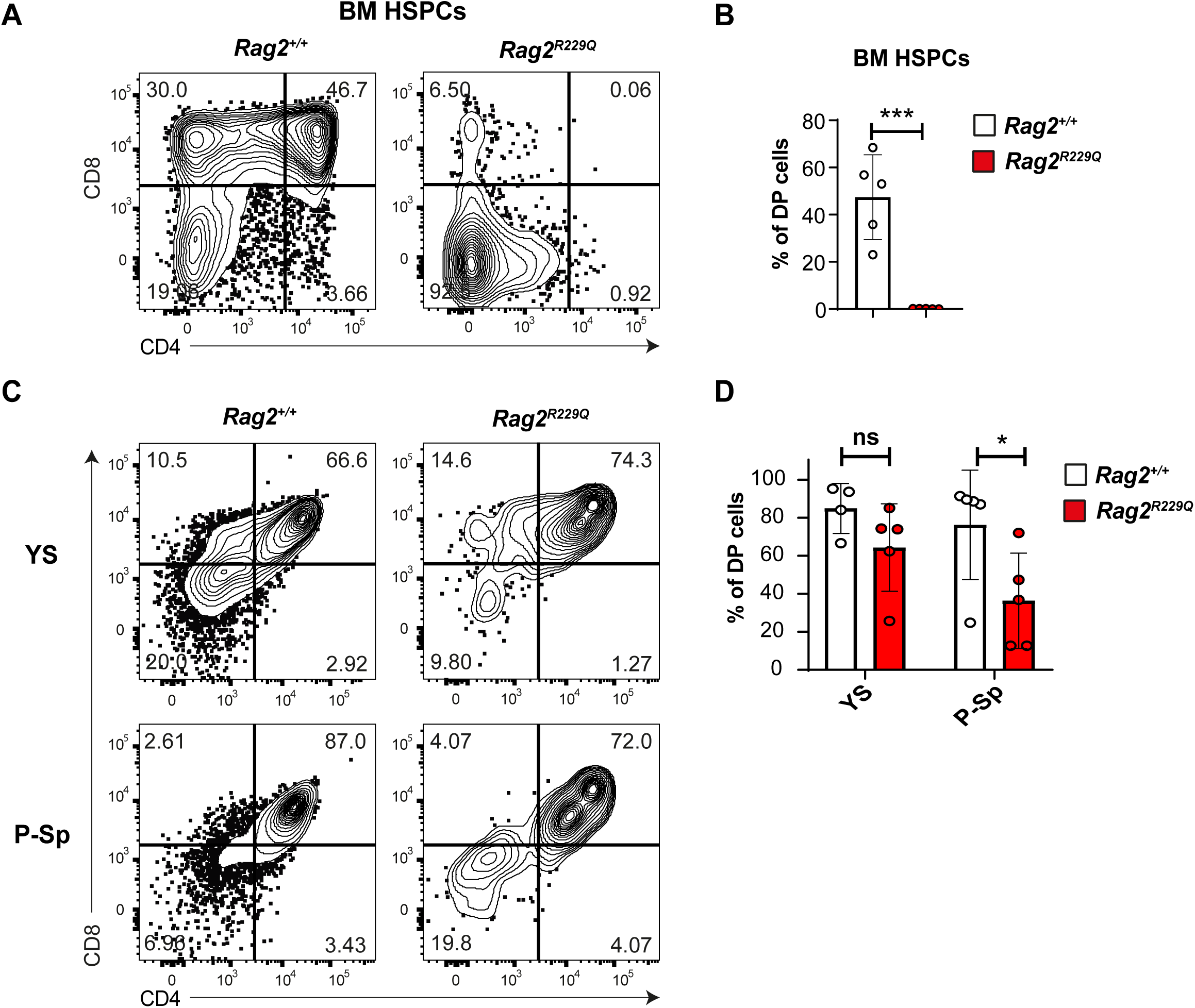
Embryonic but not adult *Rag2^R229Q^* hematopoietic progenitors can generate CD4+CD8+ double positive T-cells *ex vivo*. (A) Representative flow cytometric analysis of CD4^+^CD8^+^ T-cell potential of adult LSK. Hematopoietic progenitors were isolated from the BM of 12-week-old *Rag2^+/+^*and *Rag2^R229Q^* mice and cultured on OP9DLL1 for T-cell differentiation, gated on FSC/SSC/CD45^+^ (gating strategy shown in Figure S5E). Frequencies are shown in figure (B). n=5, independent. (C) Representative flow cytometric analysis of CD4^+^CD8^+^ T-cell potential of cells isolated from E9.5 *Rag2^+/+^* and *Rag2^R229Q^* YS and P-Sp upon culture on OP9-DLL1 for T-cell differentiation, gated on FSC/SSC/CD45^+^/CD25^neg^/CD44^neg^ DN4 cells, as shown in Supp. Fig. 2B. (D) Frequency of DP cells generated from YS and P-SP in *Rag2^+/+^*(white) and *Rag2^R229Q^* (red) cultures. n=4 *Rag2^+/+^* and n=5 *Rag2^R229Q^*embryos from 3 independent litters. p-values are for unpaired Student’s t-test. *p<0.05. ns, not significant.

### Embryonic reprogramming of *Rag2^R229Q^* BM HSPCs rescues their T-cell potential *ex vivo*

Our *ex vivo* data indicate that the *Rag2^R229Q^* OS mutation selectively affect the T-lymphopoiesis of BM-derived HSPCs. It has been shown that adult BM HSPCs can display embryonic-specific lymphoid lineage potential (e.g. B1 B-cells, Vγ3 T-cells) upon the overexpression of Lin28, a master regulator of pre-natal identity in hematopoietic cells^29^. Thus, we hypothesized that adult *Rag2^R229Q^* HSPCs should overcome their major block in T-lymphoid development once “reprogrammed” to an embryonic state. Hence, we tested whether the embryonic Lin28-induced reprogramming could rescue the lymphopoietic output of *Rag2^R229Q^* adult BM HSPCs. We designed a lentiviral vector expressing either the *Lin28* or the *GFP* (used as negative control, CTR) genes under the ubiquitous phosphoglycerate kinase (PGK) promoter with an NGFR (nerve growth factor receptor) reporter cassette to track the transduced cells. Lin^neg^ HSPCs isolated from 12-14 weeks old *Rag2^R229Q^* or *Rag2^+/+^* mice were transduced 16 hours before seeding them on OP9-DLL1 stroma for *in vitro* T-cell differentiation (Figure S3A). Upon transduction, cells efficiently expressed LIN28 (Figure S3B), whose functionality was confirmed by the downregulation of the *Let7-g* miRNA (Figure S3C), a known target of LIN28^38^. In line with the previous results, *Rag2^R229Q^* HSPCs transduced with the CTR vector did not generate DP cells *ex vivo* (Figures 3A and 3B). In sharp contrast, LIN28 expression in *Rag2^R229Q^* rescued the ability to give rise to DP (Figures 3A and 3B), and CD3^+^ cells (Figures S3D and S3E). The sorting of *Rag2^R229Q^* DP rescued cells from end stage cultures confirmed they carried the Rag2 hypomorphic mutation via genotype (Figure S3F). This result further validates our hypothesis that the ontogeny dictates the susceptibility of (DP)-lymphopoiesis to the *Rag2^R229Q^* mutation.

**Figure 3.**
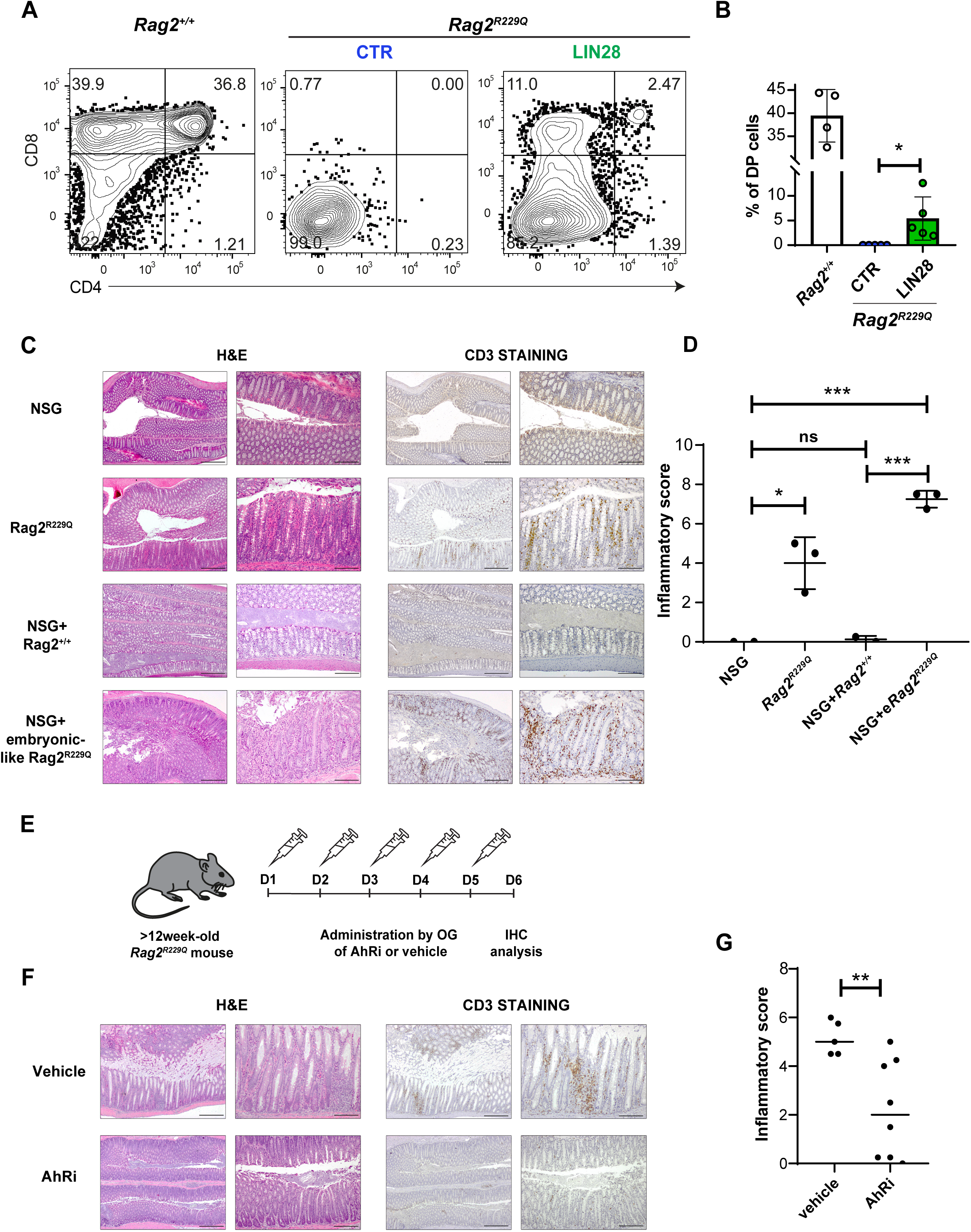
Contribution of embryonic lymphocytes to OS severe autoimmunity. (A) Flow cytometric analysis of Lin^neg^ hematopoietic progenitors isolated from the BM of 12-week-old *Rag2^+/+^* and *Rag2^R229Q^*mice, transduced overnight with CTR or LIN28 lentiviral vectors and cultured on OP9-DLL1 for T-cell differentiation. Quantifications of different experiments are shown in panel (B). n=5, independent. p-values are for unpaired Student’s t-test. *p<0.05. (C) Photo-micrographs of Hematoxylin Eosin (H&E, left panels) and CD3 (right panels) stainings of large intestine of newborn (post-natal day 3, P3) NSG immunodeficient mouse recipients of *Rag2*^+/+^ and *Rag2^R229Q^* embryonic-like T-cell progenitors. (D) Inflammatory score (as defined in method section) of NSG immunodeficient mice transplanted with *Rag2^+/+^* and *Rag2^R229Q^*embryonic-like T-cell progenitors, calculated based on combined morphological alterations and lymphoid infiltrate parameters, as detailed in the Methods section. p-values are for unpaired Student’s t-test. ***p<0.001. ns, not significant. Each dot represents a mouse. NSG, n=2; *Rag2^R229Q^*, n=3; NSG+*Rag2*^+/+^, n=2; NSG+e*Rag2^R229Q^*, n=3; (E) Schematics of AhR inhibitor (AhRi) *in vivo* administration: 12-week-old *Rag2^R229Q^* mice were treated daily by oral gavage with AhRi CH-223191 for five consecutive days and euthanized for the immunohistochemistry analysis on the sixth day. (F) Photo-micrographs of Hematoxylin Eosin (H&E, left panels) and CD3 (right panels) stainings of large intestine of 12-week-old *Rag2^R229Q^* mice treated by oral gavage with either vehicle (corn oil) or AhR inhibitor (CH-223191) for 5 consecutive days. (G) Computed inflammatory score of 12-week-old *Rag2^R229Q^*mice treated either vehicle or AhR inhibitor. n=8 *Rag2^R229Q^* treated mice from 2 independent experiments. p-values are for unpaired Student’s t-test. **p<0.01.

### Adoptive transfer of Rag2^R229Q^ embryonic-like progenitors triggers severe inflammation in immunodeficient hosts

While our results suggest that embryonic lymphocytes may serve as an alternative and/or complementary source of the residual T-cells in OS mice, their specific contribution to the OS autoimmune inflammatory phenotype has not been addressed yet. Since the Lin28-mediated embryonic reprogramming of adult BM HSPC yields a high number of cells suitable for *in vivo* adoptive transfer experiments, we assesed the ability of Lin28-transduced embryonic-like *Rag2^R229Q^* lymphoid progenitors to colonize peripheral tissues and trigger inflammation *in vivo*. For this, T-cell progenitors were differentiated from *Rag2^R229Q^* BM HSPCs over-expressing LIN28 on OP9-DLL1 stroma for 14 days, isolated by fluorescence activated cell sorting (FACS) and transplanted into the liver of newborn NOD.Cg-*Prkdc^scid^ Il2rg^tm1Wjl^*/SzJ (NSG) mice (Figure S3G). At 12 weeks post-transplant, NSG mice were euthanized, and the large intestine (the site with the major inflammatory phenotype in *Rag2^R229Q^* mice in our housing conditions) was collected and processed for immunohistochemistry analyses. Uninjected NSGs were used as negative control for peripheral lymphoid infiltrates and inflammation, while NSG mice injected with *Rag2*^+/+^ T-cell progenitors as positive control for transplantation and lymphoid peripheral repopulation. *Rag2^R229Q^* mice were used as baseline control for peripheral inflammation (Figure **3C**, 3D **and** S3H). In NSG uninjected control mice, no CD3^+^ T-cells were detected in the lamina propria, in line with the immunodeficient status of these recipients (Figure 3C). As expected, a physiological distribution of CD3*^+^* lymphocytes was observed in the intestine of NSG recipients transplanted with *Rag2*^+/+^ T-progenitors, with a regular morphology of the tissue, comparable to the one of the not injected NSG (Figure 3C). In both cases, inflammation was absent in the peripheral tissue (Figure 3D, S3H). In contrast, NSG mice transplanted with *Rag2^R229Q^*progenitors showed numerous foci of CD3^+^ T-cells infiltration in the lamina propria, in correlation with morphological alterations of the tissue architecture (Figure 3C). The grade of inflammation in the large intestine of *Rag2^R229Q^*-transplanted NSG mice was comparable to the one of the *Rag2^R229Q^* original murine model (Figure 3D, S3H). Thus, with these *in vivo* experiments, we provide strong evidence that embryonic-like *Rag2^R229Q^* progenitors can contribute to the pool of the residual OS T-cells responsible for peripheral inflammation.

### Aryl hydrocarbon receptor antagonist decreases T-cell infiltrates and ameliorates the inflammatory phenotype in the large intestine of *Rag2^R229Q^* mice

Given that pre-natal and post-natal lymphocytes development and function are regulated differently^39–42^, we reasoned that it should be possible to exploit specific vulnerabilities of embryonic lymphocytes for OS therapeutic intervention. To test this hypothesis, we evaluated the effect of the inhibition of Aryl Hydrocarbon Receptor (AhR), a crucial regulator of the homeostasis and activation of specific subsets of intra-epithelial lymphocytes, in particular those with an embryonic origin, such as Vψ3^+^ T-cells^37,43^. We treated adult *Rag2^R229Q^* mice with either the AhR antagonist CH-223191, or the empty vehicle (corn oil) for 5 consecutive days, and on the 6^th^ day we analyzed their large intestine (Figure 3E). Histological analysis showed that CH-223191 treatment resulted in an alleviation of the severe and diffuse morphological alterations, consisting of abscesses and elongated villi enriched of inflammatory cells normally observed in control animals, as well as a reduction of CD3^+^ T-cell clusters in the large intestine of most of the mice treated with CH-223191 compound (Figure 3F). As a result, CH-223191 significantly reduced the overall inflammation score (Figure 3G). These results suggest that AhR inhibitors could be an effective treatment for autoimmune manifestation in OS patients, probably targeting the activation of peripheral lymphocytes of embryonic origin.

### *Rag2^R229Q^* embryonic *ex vivo* T-cell potential is restricted to HSC-independent hematopoietic progenitors

Our preliminary data support the contribution of E9.5 YS and P-Sp to the pool of OS T-cells. Although at this stage hematopoietic progenitors are mostly HSC-independent, we cannot completely exclude the presence of early emerging HSC-precursors^44^, potentially contributing to *Rag2^R229Q^* embryonic lymphopoiesis as well. On the other hand, at E14.5 it is possible to discriminate HSC-dependent and HSC-independent progenitors that are concomitantly present in the FL, based on their marker’s expression. In fact, while HSCs segregate to the Lin^neg^Sca1^+^cKit^+^ (LSK) CD48^neg^CD150^+^ population, only the LSK CD150^neg^ cell fraction includes HSC-independent progenitors able to contribute to embryonic-restricted lineages^45^. To dissect the distinct contribution of embryonic HSCs and HSC-independent progenitos to the pool of *Rag2^R229Q^*T-cells, we isolated these fractions from E14.5 *Rag2*^+/+^ and *Rag2^R229Q^* LSK FL and tested their T-lymphoid potential *ex vivo* (Figure 4A). While both *Rag2*^+/+^ HSC-independent progenitors and HSCs gave rise to DP (Figure 4B) and CD3^+^ T-cells (Figure S4A), only the HSC-independent *Rag2^R229Q^*progenitors showed residual DP/CD3^+^ T-cell potential (Figures 4B, 4C and S4A), as *Rag2^R229Q^* HSCs failed to progress beyond the DN3 stage of T-cell development (Figure S4B). Collectively, our results strongly support a model where *Rag2*^R229Q^ OS mutation selectively affect definitive HSC-derived T-lymphopoiesis which is severely blocked at DN3 stage of *in vitro* maturation, while it allows the maturation of embryonic HSC-independent progenitors along the T-lymphoid lineage.

**Figure 4.**
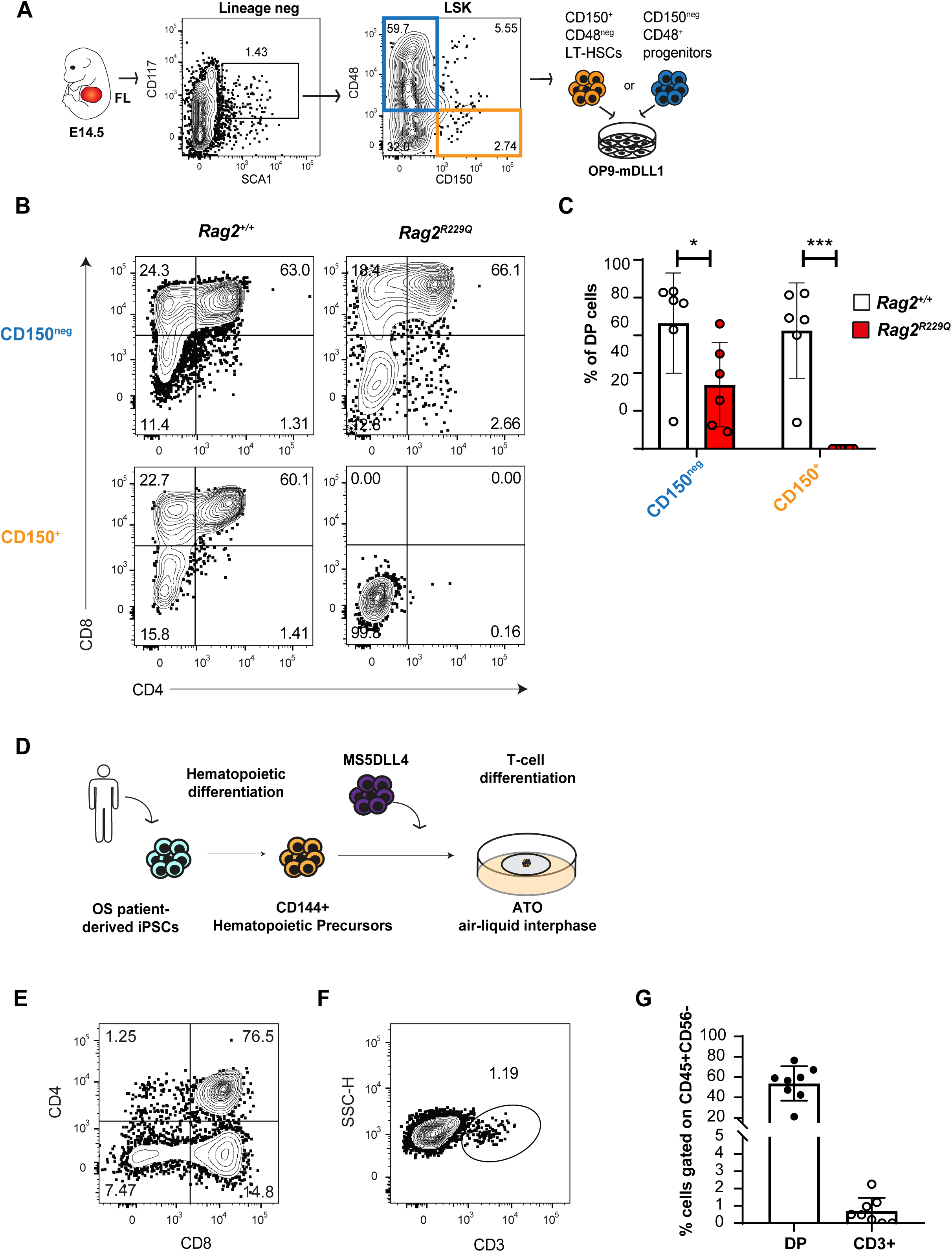
HSC-independent progenitors display T-cell potential in both *Rag2^R229Q^* mice and OS iPSCs. (A) FAC-sorting strategy to isolate long-term (LT)-HSCs (orange, LSK CD48^neg^CD150^+^) and HSC-independent progenitors (blue, LSK CD48^+^CD150^neg^) from the FL of E14.5 *Rag2^+/+^* and *Rag2^R229Q^* embryos. Both cell fractions were cultured on OP9-DLL1 for T-cell differentiation. (B) Representative flow cytometric analysis of DP T-cell potential of each fraction, gated on CD45**^+^**CD25^neg^CD44^neg^ cells (gating strategy shown in Figure S5F). (C) Frequency of DP cells in *Rag2^+/+^* (white) and *Rag2^R229Q^*(red) cultures, generated from LSK CD150^+^CD48^neg^ and LSK CD150^neg^CD48^+^ FL progenitors. Each dot represents a different embryo. n=6 *Rag2^R229Q^* embryos from 3 independent litters. p-values are for unpaired Student’s t-test. *p<0.05. ***p>0.001. (D) Schematic of T-cell differentiation of *RAG2^T125R^* OS iPSCs. (E) Representative flow cytometric analysis of DP and CD3^+^ (F) T-cell potential of *RAG2^T125R^* OS iPSCs. Gated on SSC/FSC/CD45+. (G) Frequency of DP and CD3+ T-cells generated from *RAG2^T125R^* OS iPSCs. n=8, independent.

### *In vitro* T-lymphopoiesis of iPSC-derived HSC-independent hematopoietic progenitors is compatible with OS mutations

Although animal models serve as an important tool for elucidating mechanisms of OS pathogenesis, there is a clear need to study OS in a human context. Many differences exist between human and murine T-cell development^46^, thus disease mechanisms might differ as well. However, the rarity of the disease and the poor accessibility of appropriate human tissue samples prevent the detailed cellular and molecular study of the mechanisms at the basis of OS onset. In these regards, patient-derived iPSCs give an unprecedented opportunity to answer fundamental questions related to disease pathogenesis. For this work, we used an iPSC line generated from an OS patient carrying a *RAG2* mutation leading to a C-terminus truncated protein, completely lacking the regulatory region of the protein (c.374_375delCA, p.T125Rfs*9; p.T125Rfs*9). In this OS patient, this mutation severely impairs TCR rearrangement, as assessed by the complete lack of T-cell receptor excision circles (TREC) usually detected in the young individuals, but residual CD3^+^ lymphocytes are present in peripheral tissues^47^. Using the hPSC hematopoietic differentiation protocol that we previously developed to recapitulate HSC-independent hematopoiesis^48–51^, we investigated whether embryonic T-lymphopoiesis is unaffected by OS mutations also in a human setting. CD144^+^ cells containing hematopoietic precursors were isolated at day 9 of iPSC differentiation and further cultured in artificial thymic organoids (ATO) which support T-cell development *in vitro*^52^ (Figure 4D). As expected from human hematopoietic progenitors carrying RAG mutaitions^53–55^, after 30 days of co-culture most of CD45^+^ progenitors efficiently progressed to the CD4^+^CD8^+^ DP stage (Figure 4E). Remarkably, HSC-independent OS cells can also generate CD3^+^ T-cells (Figures 4F and 4G), which are not observed from RAG-null mutants^53–55^, indicating that human OS HSC-independent hematopoietic progenitors display the ability to progress to mature stages of T-cell development *in vitro*. These results suggest that embryonic lymphocytes might contribute to OS pathogenesis also in humans and as such represent a novel therapeutic target to mitigate autoimmune manifestations in OS patients.

## Discussion

Autoimmune disorders arise from deregulated self-reactive immune response. Currently, little is known about molecular and cellular triggers of many forms of autoimmunity, thus hampering the design of targeted treatments. In this study, we sought to characterize potential cells of origin of self-reactive immunity exploiting the context of OS, a genetic disease that offers an unparalleled platform to study T-cell-mediated peripheral autoimmune-like manifestions in the absence of a functional adaptive immune system^2^. Using the *Rag2^R229Q^* mouse model, we provide evidence of the previously unappreciated contribution of embryonic lymphocytes to the pool of autoreactive T-cells. In fact, we show that *Rag2^R229Q^* embryonic HSC-independent progenitors generate CD3+ T-cells, while adult HSPCs are unable to do so, unless they are reprogrammed to an embryonic-like state via LIN28 overexpression. Remarkably, using adoptive transfer in immunodeficient recipients we further showed that these embryonic-like lymphoid progenitors can trigger autoimmune-like manifestations cell-autonomously, at least partially via the ligand activation of the AhR pathway. Our data support a model in which HSC-independent embryonic lymphocytes contribute to the inflammatory and autoimmune-like manifestations in OS patients. Given that their development is not profoundly impaired by OS mutations, these embryonic lymphocytes colonize prenatally peripheral organs, in particular those at the interface with the external environment, where they represent the first line of defense protecting the host. In contrast, OS mutations severely impair the development of HSC-dependent lymphopoiesis, including the correct establishment of central and peripheral tolerance^56,57^.

As such, the lack of functional regulatory T-cells (Treg) favors the abnormal expansion of these self-responding embryonic T-cell clones, resulting in heightened immune activation and increased vulnerability to epithelial damage^30^. This model is supported by the observation that residual lymphocytes present in OS patients display characteristics that are proper of the very first lymphocytes generated in the embryo before the onset of HSCs, including a highly restricted TCR repertoire that is shared across patients and is often specific for the recognition of self-antigen^58^. In addition, the average length of complementary diversity region (CDR)3 in human OS autoreactive peripheral lymphocytes is shorter compared to healthy samples^11^. Interestingly, TdT enzyme, which is responsible for TCR/BCR diversity and CDR3 elongation, is not expressed at the first stages of embryonic development^59–61^. Thus, it is reasonable to think that OS lymphocytes with short CDR3 have been generated in the early embryo.

However, the model we are proposing does not completely exclude the contribution of adult HSC-derived T-lymphocytes to OS autoimmunity. Indeed, the expanded and autoreactive OS clonotypes that are present in the peripheral tissues could also derive from rare thymic progenitors or from immature DN T-cells undergoing final maturation and antigen-mediated expansion in peripheral tissues, therefore after thymic egression. As such, OS T-cell repertoire might be composed by T-cells coming from both embryonic DP thymocytes and adult DN thymocytes, accomplishing maturation and expansion in peripheral tissues. However, all our results strongly suggest that embryonic T-lymphopoiesis is less susceptible than the adult counterpart to hypomorphic RAG2 mutations. Overall, this study describes the contribution of pre-natal lymphocytes to genetic forms of autoimmunity, specifically in the case of OS. The embryonic ontogeny of T-cells displaying a deregulated activation could be explored in other human autoimmune disorders, such as Type 1 Diabetes, Systemic Sclerosis or Rheumatoid Arthritis, similarly characterized by either reduced thymic output, shortened and hypovariable CDRs and/or oligoclonal expansion of auto-reactive lymphocytes triggering inflammation and tissue damages^62–65^. As such, this study establishes a new paradigm for the origin of autoimmune manifestations and paves the way for the design of better and more targeted treatments for these types of disorders.

## Acknowledgements

We thank A.D. and A.V. lab members for inputs and critical reading of the manuscript; Mario Squadrito for the help in the viral vectors’ preparation and the Flow cytometry Resource, Advanced Cytometry Technical Applications Laboratory (FRACTAL) at Ospedale San Raffaele. Tom Taghon for providing the MS5-DLL4 stromal cell line. L.N. was supported by the Division of Intramural Research, National Institute of Allergy and Infectious Diseases, NIH (grant AI001222). A.V. was supported by the EU H2020 grant RECOMB(755170-2). A.D. was supported by the Telethon Foundation (TGT16C04 and TGT16G03B). S.C. conducted this study as fulfillment of an international Ph.D. in Molecular Medicine, Vita-Salute San Raffaele University. Animal studies and human pluripotent stem cell studies have been approved by the Italian Ministry of Research (IACUC 1092) and by the Ospedale San Raffaele Ethical Committee (TIGET-HPCT), respectively.

## Author Contributions

A.D. formulated the initial concept. S.C., A.V. and A.D. designed the experiments and analyzed the data; S.C., R.S., R.R., V.C., E.D. and A.D. performed the experiments; E.F. performed all the immunohistochemistry analyses; L.N. and K.D. generated the OS iPSC line; S.C. and A.D. wrote the manuscript. S.C., A.D., A.V., V.C., R.S. and E.F. revised the manuscript.

## Methods

### Mouse strains

The C57/Bl6 *RAG2^R229Q/R229Q^*OS mouse model was originally established by Dr. Anna Villa’s research group at San Raffaele Telethon Institute for Gene Therapy (SR-TIGET)^30^. Mice were maintained in heterozygosis (RAG2^+/R229Q^) which display a phenotype indistinguishable from *Rag2^+/+^*littermates. We intercrossed RAG2^+/R229Q^ mice to produce homozygous *Rag2^R229Q/R229Q^* experimental mice, which display an OS-like phenotype. NOD.Cg-*Prkdc^scid^ Il2rg^tm1Wjl^*/SzJ (NSG) mice were bred and newborn mice at post-natal day3 (P3) were used as recipients for intra-liver transplant of RAG2^+/+^ and *Rag2^R229Q^* T-cell progenitors. All animal procedures were performed according to protocols approved by the Animal Care and Use Committee of the Ospedale San Raffaele (IACUC 1092) and communicated to the Ministry of Health and local authorities according to the Italian law.

### Analysis of T-cells and B-cells in murine thymus and spleen

12–14-week-old *Rag2*^+/+^ and *Rag2^R229Q^*mice were euthanized by CO_2_ and cervical dislocation, disinfected with a 70% EtOH solution and fixed on their back on a styrofoam block for tissue collection. For processing, thymi and spleens were smashed, and the resulting cell suspension was strained through a 40μm nylon filter and washed in cold PBS (Sigma-Aldrich, D8662) +10% FBS (Hyclone, 12389802). Cells were centrifuged at 1500 rpm for 3 minutes and incubated with an anti-mouse CD16/CD32 blocking antibody for 15 minutes at 4°C. After the first incubation with the blocking reagent, cells were stained with specific anti-mouse antibodies for 15 minutes at 4°C. For T-cell progenitors’ analysis, samples were stained with anti-CD45, -CD4, -CD8 and -CD3 antibodies. For the flow cytometry analysis of splenic B-lymphocytes, we stained spleen samples with antibodies anti-CD45, -CD21, -CD23, -CD19, and -IgM. After the incubation, samples were washed and resuspended in PBS containing 2% FBS and 7-AAD for FACS analysis.

### Analysis of B1 and B2 B-cells in murine peritoneal wash

The analysis of B1 and B2 B cells was performed on peritoneal wash taken from 12–14-week-old mice, euthanized and prepared as explained above. By using forceps to uplift the skin of the abdomen, ice cold PBS containing 10% FBS was injected into the peritoneal cavity through a 27-gauge needle-syringe, while being careful not to puncture any organ. After injection, the mouse abdomen was gently massaged to dislodge any attached cell into the PBS solution. With forceps, an incision was made in the lower abdomen and cavity fluid was collected with a pipette. Cell suspension was filtered, centrifuged at 1500 rpm for 3 minutes and resuspended in PBS + 10% FBS for staining. For flow cytometry analysis of B1 and B2 lymphocytes, we stained samples with antibodies anti-CD45, -B220, -CD5, -CD19, and -IgM. After the incubation, the samples were washed and resuspended in PBS containing 2% FBS and 7-AAD for the analysis.

### Analysis of murine skin and intestine tissue-resident T-cells

For back skin collection, mice were shaved to remove as much fur as possible. The entire back skin, from shoulders to just above the tail, was cut away from adult mice with scissors and forceps and put in a dish with ice-cold PBS. Skin was then cut into small pieces (2-4mm) and weighed on a fine balance to put 0.6g of skin per 3.5ml of digestion medium (0.385mg/ml Liberase TL-Roche 246749-in HBSS/PBS). Samples were incubated at 37°C for 40-60 minutes while shaking. For the analysis of the large intestine, the intestinal tract from the anus to the caecum (excluded), was dissected with scissors and forceps, and feces were eliminated from the tissue. Large intestines were opened longitudinally and cut into 3-5 mm pieces. Tissues were washed in HBSS buffer (Corning 21-020-CV) (0.5M EDTA, 1mM DTT, 15mM Hepes and 10% FBS) under agitation and then digested at 37°C for 30 minutes with Liberase (0.15mg/ml Liberase TL-Roche 246749-in HBSS/PBS). For both skin and intestine, the liberase-enzymatic reaction was blocked by adding RPMI (Corning 10-040-CV)+10%FBS, and the digested cell suspension was strained with a 70µm filter. Before staining, cells were incubated with an anti-mouse CD16/CD32 blocking antibody for 15 minutes at 4°C. Then, cells were stained with anti-mouse antibodies for 15 min at 4°C. For the analysis of skin and intestinal T-lymphocytes, we stained samples with antibodies anti-CD45, -CD3, -TCRβ, and -TCRγδ. After the incubation with primary antibodies, samples were washed and resuspended in PBS containing 2% FBS and 7-AAD for the FACS analysis.

### Bone Marrow isolation

12–14-week-old *Rag2*^+/+^ and *Rag2^R229Q^*mice were euthanized by CO_2_, mounted on the Styrofoam block and bones from hips to ankles were dissected and put in a Petri dish with ice cold PBS. Any remaining muscles or connective tissues attached tothe bones were removed by using scissors and forceps. For the isolation of bone marrow, bones were transferred to a mortar with PBS 10% FBS and crushed by using a pestel. Cell suspension was filtered through a 70µm filter, collected into a falcon tube and centrifuge at 1000 rpm for 5 min. Lin^neg^ cells were isolated through negative magnetic bead selection with Lineage negative depletion kit, according to manufacturer’s instructions (Miltenyi Biotech, 130-090-858) and used for experiments.

### Embryonic tissues dissection and isolation

For these experiments, heterozygous *Rag2^+/R229Q^* mice were bred and embryonic development was estimated considering the day of vaginal plug formation as 0.5 day post-coitum (E 0.5). At E9.5 and E14.5, pregnant mice were euthanized for the retrieval of embryos. The dissection procedure was performed in a horizontal laminar flow hood with a stereomicroscope. The uterus was transferred into a Petri dish with PBS and cut into single embryo units with scissors and forceps. The outer muscular membranes of the uterus and the decidua were dissected away to expose the YS and the embryo proper. First, YS was gently separated from the embryo proper with a net cut at the origin of the vitelline vessel. For P-Sp isolation, the caudal part was separated from the rest of the embryo and somites were dissected away by using fine needle-syringes. In E14.5 embryos, the FL was isolated from the rest of the embryo body by needle-syringes. The “tail” of the embryos was used for genotypic analysis. For tissue digestion, samples were treated with 1mg/ml Collagenase Type I (Sigma Aldrich SCR103) in water-bath at 37°C for 15 minutes and triturated by pipetting up and down several times. Enzymatic reaction was blocked by adding PBS+10% FBS+1% DNAse (Calbiochem, 260913) and cell suspension was filtered and centrifuged at 1000 rpm for 5 min. At that point, the cell pellet was ready to be plated or eventually sorted for the for the T-cell *in vitro* assay.

### Cloning and lentiviral vectors preparation

Lin28 coding sequences (CDS) have been amplified from mouse ESC cDNA by Phusion High Fidelity PCR, according to manufacturer instructions (New England Biolab, M0530S). Primers (Lin28_Fw_AgeI:AAAAAACCGGTCTTTGCCTCCGGACTTCTCTGG; Lin28_Rv_SalI:AAAAAGTCGACAAAGACAGGGTGACACTGGGA) have been designed to obtain compatible ends for the insertion of the CDS into the hPGK.GFP.mhCMV.dNGFR.SV40PA.WPRE plasmid, at the site of GFP protein. Purified PCR products and the plasmid backbone were digested with *Age*I/*Sal*I restriction enzymes, purified, mixed in a ratio 1:3 (insert:backbone plasmid) and ligated with T4 ligase according to manufacturer’s instructions (Quick Ligation kit, NEB M2200). The ligation reaction was transformed into competent TOP-10 bacteria and plated on agarose plates with Ampicillin. Single transformed ampicillin resistant colonies were grown in LB + Ampicillin and their DNA was extracted by the Wizard® Plus SV Minipreps DNA Purification System kit (Promega, A1460). Clones with correct insertions were evaluated by Sanger sequencing (LightRun tube, Eurofins service). Positive cones were amplified, and a maxi-prep was performed for proceeding to vector production (Wizard® PlusMaxipreps DNA Purification System N2511). Third generation lentiviral (LV) vectors have been prepared and tittered as described in literature (Dull et al, 1998; Follenzi et al, 2000). Briefly, self-inactivating (SIN) LV vectors were produced using the transfer vectors, the packaging plasmid pMDLg/pRRE, Rev-expressing pCMV-Rev, the pADV and the VSV-g envelope-encoding pMD2.VSV-G plasmids. The resulting bidirectional LVs have been used to overexpress the CDS of candidate murine genes Lin28 or eGFP (as control) under the control of the phosphoglycerate kinase (PGK) promoter and the NGFR and the minimal cytomegalovirus (mCMV) promoter forming the antisense expression unit.

### Lin28-induced fetal reprogramming of adult HSPCs

For these experiments, Lin^neg^ cells have been FACS-sorted by FACSAria Fusion from the bone marrow of adult mice, isolated as described above. After sorting, cells were plated in StemPro™-34 SFM (Thermo Fisher Scientific 10639011), supplemented with 2% Glutamine, 1% of Penicillin/Streptomycin, 20ng/ml mTPO, 50ng/ml mSCF, 10ng/ml mFLT3L, 10ng/ml mIL3, 20ng/ml mIL6 (all the cytokines are from Miltenyi Biotech), into 96/48 multiwell plates, according to the cell number. Lin^neg^ cells were transduced with the NGFR reporter lentiviral vectors prepared as above and 16 hours later, seeded on OP9-DLL1 for T-cell differentiation.

### RNA extraction, reverse transcriptase (RT)-PCR and quantitative real-time (q)(RT)-PCR

RNA was isolated using the SV Total RNA Isolation System kit (Promega, Z3105). RT-PCR for mRNA transcripts was performed by using the ImProm-II Reverse Transcription System kit (Promega A3800). Real Time PCR was performed with Applied Biosystems Fast SYBR Green Master Mix (Thermo Fisher Scientific, 43-856-12). RNA for microRNA expression analysis was isolated using the miRNeasy Micro Kit (Qiagen 1071023). The extracted RNA was retro-transcribed using the miScript II RT kit (QIagen, 218161). qPCR for miR quantification was performed with miScript SYBR Green PCR kit (QIagen, 218073). qRT-PCR was run for 40 cycles using the Viia 7 instrument and Viia 7 software was used to extract the raw data (Ct). To determine gene expression, the difference (ΔCt) between the threshold cycle (Ct) of each gene and that of the reference gene was calculated by applying an equal threshold. Relative quantification values are calculated as the expression of the gene of interest over the expression of the reference in the same sample, by the formula 2^-ΔCt^. Let-7g miR analysis was performed by using Mm_Let-7g_2 miScript Primer Assay (Qiagen, MS00010983) and RNU6-2 (Hs_RNU6-2_11 miScript Primer Assay, Qiagen, MS00033740) as housekeeping miRNA.

### Protein extraction and WB analysis

Protein extraction was performed by resuspending cell pellets in RIPA-NP40 Buffer (150mM NaCl, 1% NP40, 0.1% SDS, 50mM TrisHCl pH 7.4, 0.5% deoxycholate) with protease inhibitor, vortexing and incubating it on ice for 10 minutes. After centrifugation at maximum speed for 2 minutes, supernatant containing protein suspension was collected and quantified with Bradford reagent (Sigma, B6916-500ML) at the spectrophotometer. As standards, Bovine Serum Albumin (BSA) serial dilutions were prepared and quantified at the spectrophotometer. A standard amount of protein (13-20ng) was denatured in a total volume of 40µl with LDS buffer and Reducing Agent (ThermoFisher Scientific, B0009) at 70°C for 10 min. Samples were subjected to SDS-PAGE in Bolt MES Buffer (ThermoFisher Scientific, B0002) to be transferred to PVDF membrane by electroblotting iBlot 2 Dry Blotting System. Then, the membranes were blocked with TBS-Tween+5%BSA for 2 hours at room temperature, while shaking. For the analysis of LIN28, membranes were blotted with rabbit polyclonal antibody (Ab) raised against LIN28 (1:10000 dilution, Cell Signaling #3978) resuspended in TBS-Tween+5% BSA, overnight at 4°C. Rabbit anti-Histone 3 polyclonal Ab (1:1000 dilution, from Abcam ab1791) was used as a normalizer. Samples were then washed with TBS-Tween and incubated with HorseRadish Peroxidase-linked secondary antibodies (Amersham ECL Rabbit IgG, HRP-linked F(ab’)₂ fragment, GE Healthcare NA9340-1ML) resuspended in TBS-Tween+5%BSA for one hour at room temperature on shaker. After 3 washes with TBS-Tween, the membranes were treated in the dark with the reagent “Immobilon Forte Western HRP substrate” (Sigma Aldrich, WBLUF0100) and visualized at ChemiDoc instrument (BIO RAD).

### OP9-DLL1/MS5-DLL4 culture

The MS5-DLL4 line was a kind gift from prof. Tom Taghon (Ghent University). The OP9-DLL1/MS5-DLL4 cell lines were maintained in Alpha Minimum Essential Medium (Thermo Fisher, 12000063) with 2.2 g/L sodium bicarbonate (Corning, 61-065-RO), supplemented with 10% FBS (HyClone CHA1115L), penicillin (100 IU/ml), streptomycin (100 μg/ml) and 1% glutamine. The lines were cultured on petri-dishes (d=10 cm) at 37°C with 5% CO2 and split with trypsin-EDTA every two days with a ratio 1:4, when they reach 80% of confluency.

### Murine T-cell differentiation

For testing T-cell potential, murine hematopoietic precursors were plated on confluent OP9-DLL1 coated 24-multiwell plates. Embryonic samples such as YS and P-Sp explants were plated as 1 embryo equivalent (ee)/well. Differentiating cells were cultured in Alpha Medium Minimum Essential Medium (Gibco, 12000-063) with 2.2g/L sodium bicarbonate, supplemented with 20% FBS (HyClone), penicillin (100 IU/ml), streptomycin (100μg/ml) and 1% glutamine. Base medium is enriched with SCF 50ng/ml (only for the first week) and 50ng/ml IL7, to engage T-cell lineage specification. Every 10 days, cells are collected, filtered and expanded on fresh stroma, with fresh cytokines-containing medium, and a small fraction is analyzed at each timepoint by FACS to assess the quality of differentiation.

### *In vivo* transplantation of *Rag2^+/+^* and *Rag2^R229Q^* progenitors in NSG mice

*Rag2^+/+^* or *Rag2^R229Q^*-reprogrammed BM Lin^neg^ hematopoietic progenitors were isolated and seeded onto OP9-DLL1 for T-cell differentiation, as explained above. After 14 days of differentiation, T-cell progenitors were isolated from the OP9-DLL1 co-culture by FAC-sorting of the CD45+ cells, resuspended in a volume of 30ul of T-cell medium and injected into the liver of newborns NSG at post-natal day 3, by using a syringe. Newborns NSG were sub-lethally irradiated the day before transplantation. A minimum of 2 million CD45+ cells was injected per mouse, since lower number of cells did not give a positive engraftment result at the time of the analysis. At 21 days post-injection, mice were weaned, and males and females were separated in different cages. At 12-weeks post-transplant, mice were euthanized by CO_2_ and tissues were collected for IHC analysis.

### CH-223191 administration by oral gavage

50mg of AhR inhibitor CH223191 (Tocris, cat. num. 3858) were resuspended in 2.5ml of DMSO and 22.5ml corn oil to reach a final concentration of 2mg/ml. 100ul were administered by oral gavage to each mouse, for a daily dosage of 200ug/100ul. The same volume of corn oil was administered to the vehicle-control group. *Rag2^R229Q^* adult mice (>12 weeks-old) were treated for five consecutive days and on the 6^th^ day of treatment they were euthanized for the IHC analysis of the large intestine.

### Histology

Mouse tissue samples were formalin-fixed and paraffin-embedded. Sections (1,5 µm) were used for routine hematoxylin and eosin (H&E) staining to check for basic histopathological changes. Moreover, sections were de-waxed, rehydrated, endogenous peroxidase activity blocked by 0.1% H_2_O_2_ for 15 minutes and nonspecific background reduced with Rodent Block (Biocare Medical) before microwaves antigen-retrieval treatment (EDTA buffer pH 8.0). In the end, sections were incubated for 1 hour at room temperature with the primary rabbit polyclonal antibody anti-CD3 (1:100; ThermoFisher Scientific), then incubated for 30 minutes with MACH 1™ Universal HRP Polymer Kit (Biocare Medical), and reactions were developed in Biocare’s Betazoid DAB and nuclei counterstained with Haematoxylin. Digital images were acquired by Olympus XC50 camera mounted on a BX51 microscope (Olympus), with CellF Imaging software (Soft Imaging System GmbH).

### Histological Score

Gut histological score was adopted for evaluating different manifestations: grade of inflammation (scoring from 0 to 4, where 0 is normal condition and 4 is a severe grade of inflammation), the structural changes of the glands (scoring from 0 to 3, where 0 is normal condition and 3 is a severe structural changes) and the goblet cell alterations (scoring from 0 to 3, where 0 is normal condition and 3 is a severe alterations).

### Human induced Pluripotent Stem Cells (iPSCs)

SCID 200-CH cells are hiPSCs reprogrammed by Sendai virus from fibroblasts of an OS patient, carrying a *RAG2* mutation (c.374_375delCA, p.T125Rfs*9). Skin biopsies were obtained and iPSC were generated according to the protocol # 0409113R approved by the Institutional Review Board of the Boston Childern’s Hosptial. The use of human iPSCs was approved by the Ospedale San Raffaele Ethical Committee, included in the TIGET-HPCT protocol.

Human iPSCs were grown on irradiated mouse embryonic fibroblast (MEF) feeders in hES medium defined as DMEM/F12 medium (Corning, L022046-10092CVR) supplemented with 25% of KnockOut™ Serum Replacement (Thermo Fisher Scientific, 10828028), 1% Penicillin-Streptomycin (Lonza, DE17-603E), 2 mM L-Glutamine (Lonza, BE17-605E), 0.1% β-Mercaptoethanol (Sigma Aldrich, M3148), 0.7% of MEM Non-Essential Amino Acids Solution (Thermo Fisher Scientific, 11140035). 1 µg/ml Ciprofloxacin HCl (Sigma Aldrich, PHR1044-1G) and 20 ng/ml human recombinant basic fibroblast growth factor (bFGF, R&D, 233-FB-500/CF) were added to hiPSCs medium right before usage. Cells were maintained and expanded at 37°C, 21% O2, 5% CO2.

### Hematopoietic and T-cell differentiation protocol of iPSCs

Human iPSCs were differentiated into hematopoietic progenitors as previously described^49^. Four or five days after thawing, hPSCs were expanded on new MEF for additional 4/5 days and then cultured on Matrigel-coated plasticware (Corning Life Sciences, 356230) for 24-48 hours, followed by embryoid body (EB) generation. EB aggregates were resuspended in SFD media defined as 75% IMDM (Corning, 15343531), 25% Ham’s F12 (Corning, 10-080-CVR), 0.005% BSA – Fraction V, B27 supplement (Thermo Fisher Scientific Cat # 12587010), N2 supplement (Thermo Fisher Scientific Cat # 17502048), 1% Penicillin-Streptomycin, 1 µg/ml Ciprofloxacin. The differentiation media was supplemented as previously described^49,50^. Briefly, the first day of differentiation SFD medium was supplemented with 2 mM L-glutamine, 1 mM ascorbic acid (Sigma Aldrich, A4544), 400 µM 1-Thioglycerol solution (Sigma-Aldrich, M6145), 150 μg/ml transferrin (R&D, 2914-HT), and 10 ng/ml BMP4 (R&D, 314-BP-MTO). 24 hours later, 5 ng/ml bFGF (R&D, 233-FB-500/CF) was added. At the second day of differentiation, 3 μM CHIR99021 (Cayman Chemical Company, CT99201) were added, as indicated. On the third day, embryoid bodies (EBs) were changed to StemPro-34 media (Thermo Fisher Scientific, 10639011) supplemented with Penicillin-Streptomycin, L-glutamine, ascorbic acid, 1-Thioglycerol and transferrin, as above, with additional 5 ng/ml bFGF and 15 ng/ml VEGF (R&D, MAB3572). On day 6, 10 ng/ml IL-6 (130-093-934), 25 ng/ml IGF-1 (130-093-887), 5 ng/ml IL-11 (130-103-439), 50 ng/ml SCF (130-096-696), 2 U/ml EPO (Peprotech, 100-64) were added. All cytokines were purchased from Miltenyi Biotec, unless indicated differently. All differentiation cultures were maintained at 37°C. All embryoid bodies and mesodermal aggregates were cultured in a 5% CO2, 5% O2, 90% N2^44^. For human T-cell differentiation, the 3D Artificial Thymic Organoid (ATO)^52^ system was adapted to our protocol. At day 9 of EB-based differentiation culture, CD144+ hematopoietic precursors were sorted with magnetic beads (Miltenyi Microbeads 130-097-857), according to the manufacturer’s instruction, and mixed with MS5-DLL4 cells (kind gift of Tom Taghon, University of Ghent)^66^ in a 1:50 ratio. Hematopoietic progenitors and stromal cells were centrifuged together at 1500 rpm for 5 minutes and resuspended in the T-cell differentiation medium, consisting of RPMI 1640 (Corning 10-040-CV), 4% B27 (Thermo Fisher Scientific 17504044), 30uM ascorbic acid and 1% P/S and Glutamax (Thermo Fisher #35050061). Each single aggregate was composed of at least 10000 CD144+ cells and 500000 MS5-DLL4 cells and seeded in a volume of 5ul onto the cell insert membrane for the air-liquid interphase culture. For the media change and harvesting for FACS analysis, the original protocol was followed^52^.

### Flow cytometry

All cytometric analyses were performed using the FACS Canto instrument (BD Biosciences) and analyzed with the FlowJo software. For antibodies, see the table below.

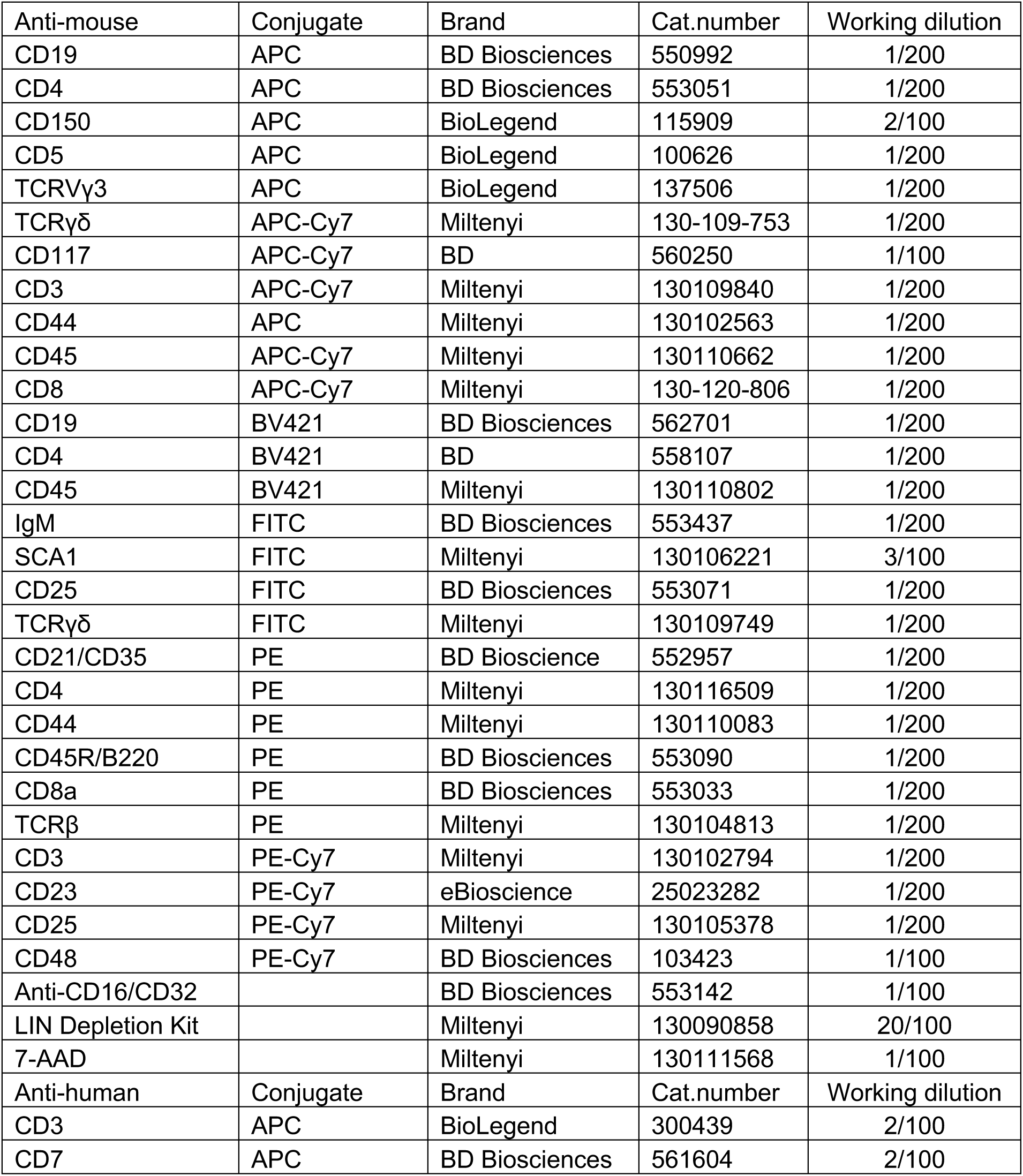
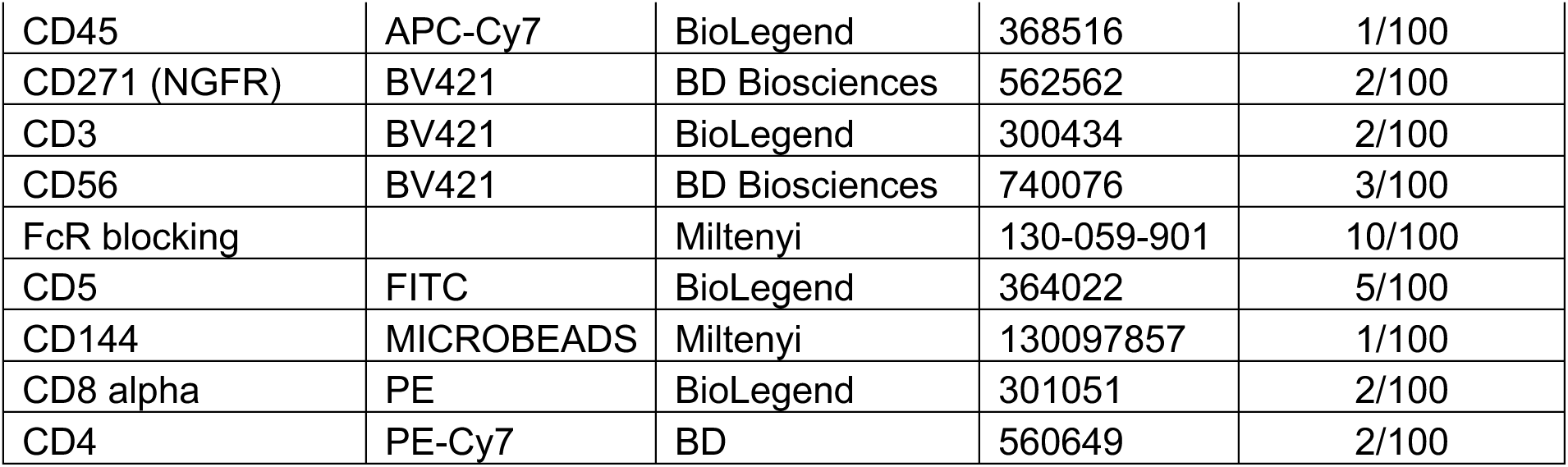

## Supplementary figure legends

**Supplementary Figure 1.**
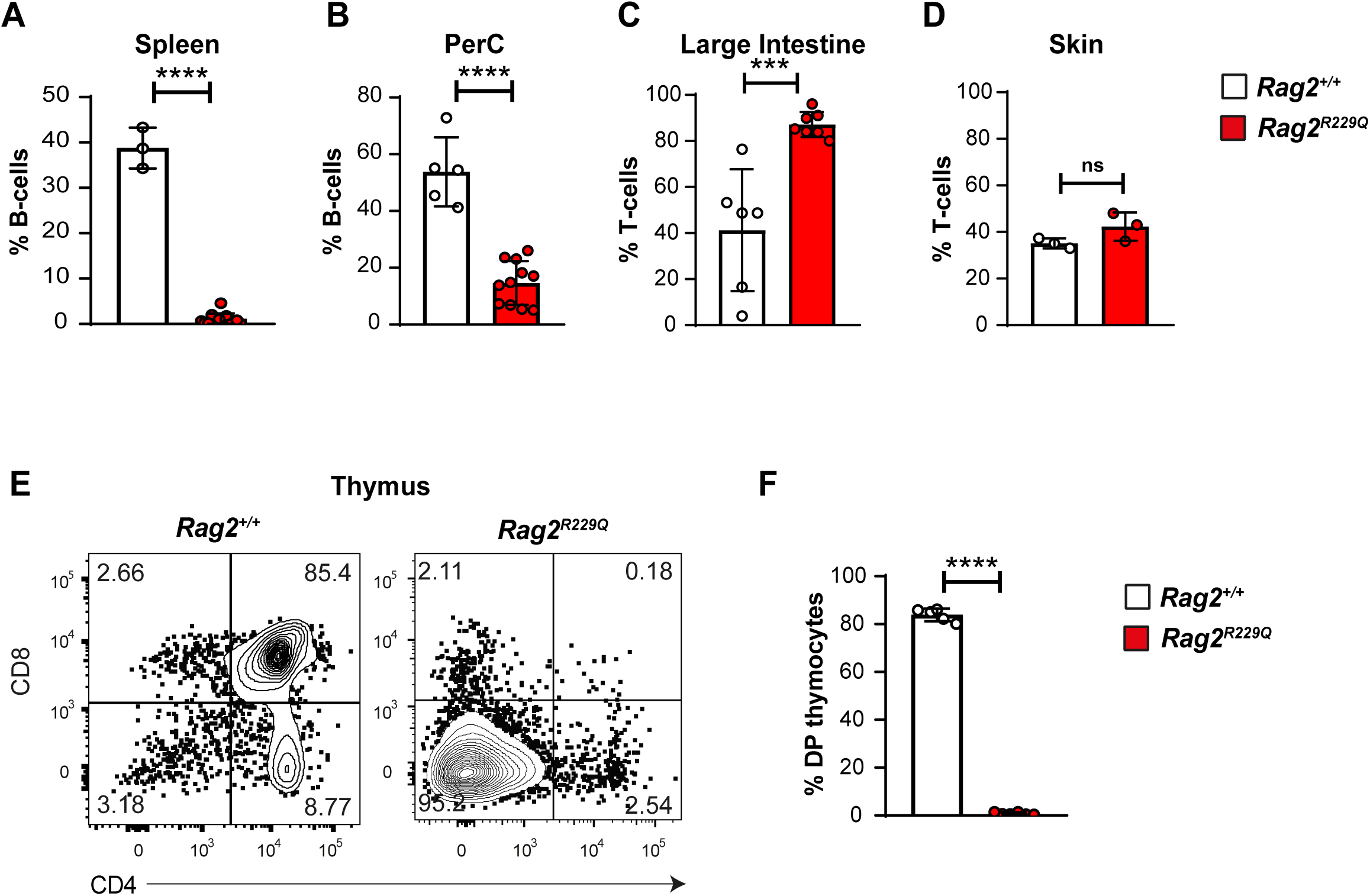
Mature lymphocytes are present in the peripheral organ of *Rag2^R229Q^* mice, but T-lymphopoiesis is impaired before the DP stage in the thymus. (A) Frequency of the splenic CD19^+^IgM^+^ B-cells on the total CD45^+^7AAD^neg^ population. n=3 *Rag2^+/+^*and n=13 *Rag2^R229Q^* mice from 3 independent litters. (B) Frequency of the peritoneal CD19^+^IgM^+^ B-cells on the total CD45^+^7AAD^neg^ population. n=3 *Rag2^+/+^* and n=13 *Rag2^R229Q^*mice from 3 independent litters. (C) Frequency of the intestinal CD3^+^ T-cells on the total CD45^+^7AAD^neg^ population. n=6 *Rag2^+/+^* and n=7 *Rag2^R229Q^*mice from 2 independent litters. (D) Frequency of the skin TCR^+^ T-cells on the total CD45^+^7AAD^neg^ population, in *Rag2^R229Q^* (red) and *Rag2^+/+^* (white) mice. n=3 *Rag2^+/+^* and n=3 *Rag2^R229Q^* mice from 2 independent litters. (E) Representative flow cytometric analysis of CD4/CD8 thymocytes in *Rag2^R229Q^* and *Rag2^+/+^* mice. Gating strategy shown in Figure S5D. Frequency is shown in (F). n=5 *Rag2^+/+^* and n=5 *Rag2^R229Q^* mice from 2 independent litters. p-values are for unpaired Student’s t-test; ***p<0.001, ****p<0.0001.

**Supplementary Figure 2.**
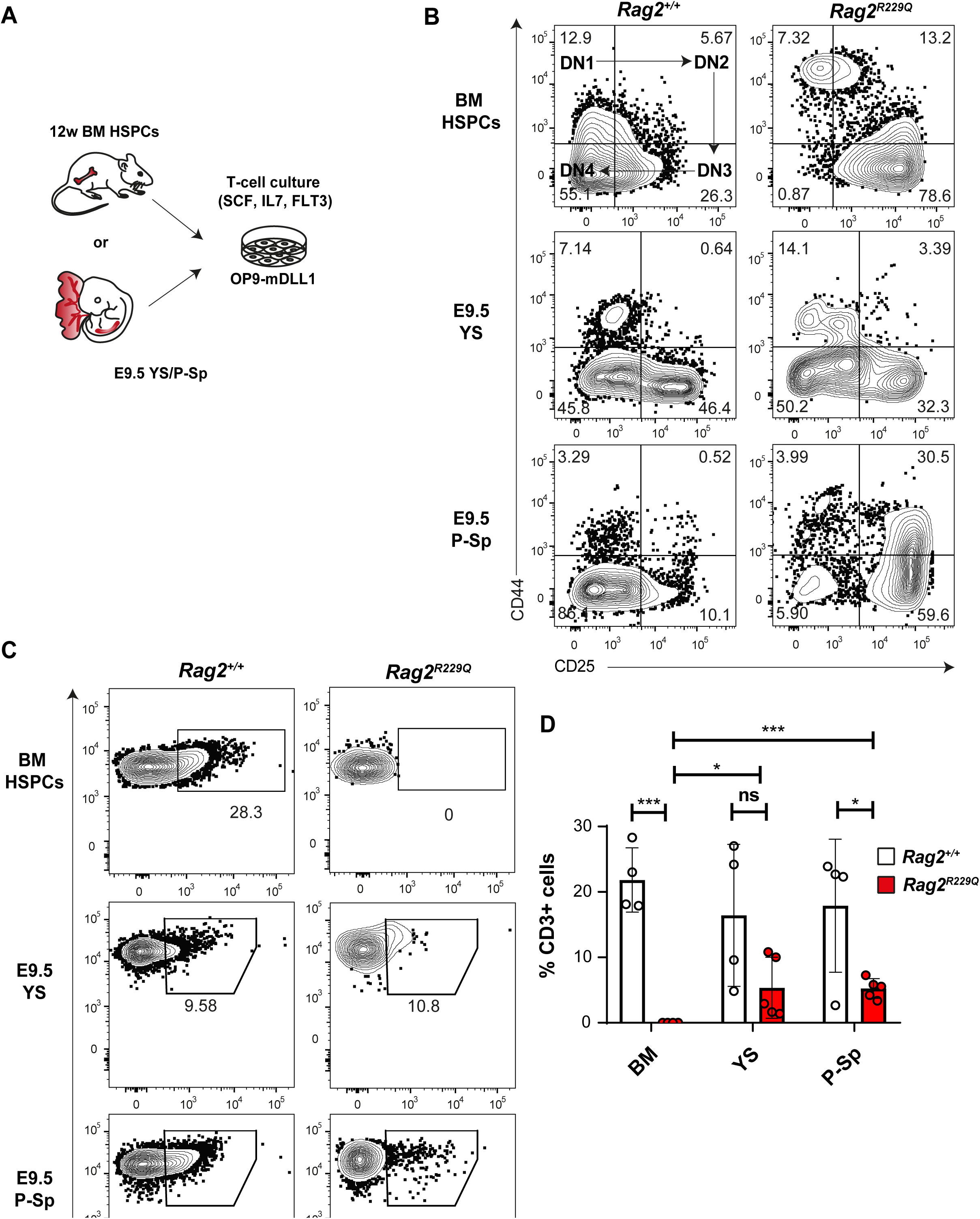
DN3-to-DN4 transition is impaired in adult BM-derived *Rag2^R229Q^* HSPCs, but not in E9.5 YS and P-Sp progenitors. (A) Experimental layout: BM HSPCs isolated from 12-weeks-old *Rag2^+/+^*and mice *Rag2^R229Q^* mice and E9.5 *Rag2^+/+^* and *Rag2^R229Q^* YS and P-Sp explants were cultured on OP9-DLL1 in T-cell differentiation medium for 30 and 20 days, respectively. (B) Rapresentative plots showing the progression of lymphopoiesis from double negative (DN) stage 1 to 4, assessed by the flow cytometric analysis of CD25 and CD44 expression (DN1:CD44^+^CD25^neg^, DN2: CD44^+^CD25^+^, DN3: CD44^neg^CD25^+^, DN4: CD44^neg^CD25^neg^) in BM HSPC, E9.5 YS and E9.5 P-Sp isolated from *Rag2^+/+^* and *Rag2^R229Q^* mice. (C) Representative flow cytometric analysis of CD3 expression in adult BM and E9.5 YS/P-Sp cultures. Cells were gated on total CD45+ cells. Quantification is shown in (D). n=4 *Rag2^+/+^* and n=5 *Rag2^R229Q^* embryos from 3 independent litters. p-values are for unpaired Student’s t-test.

**Supplementary Figure 3.**
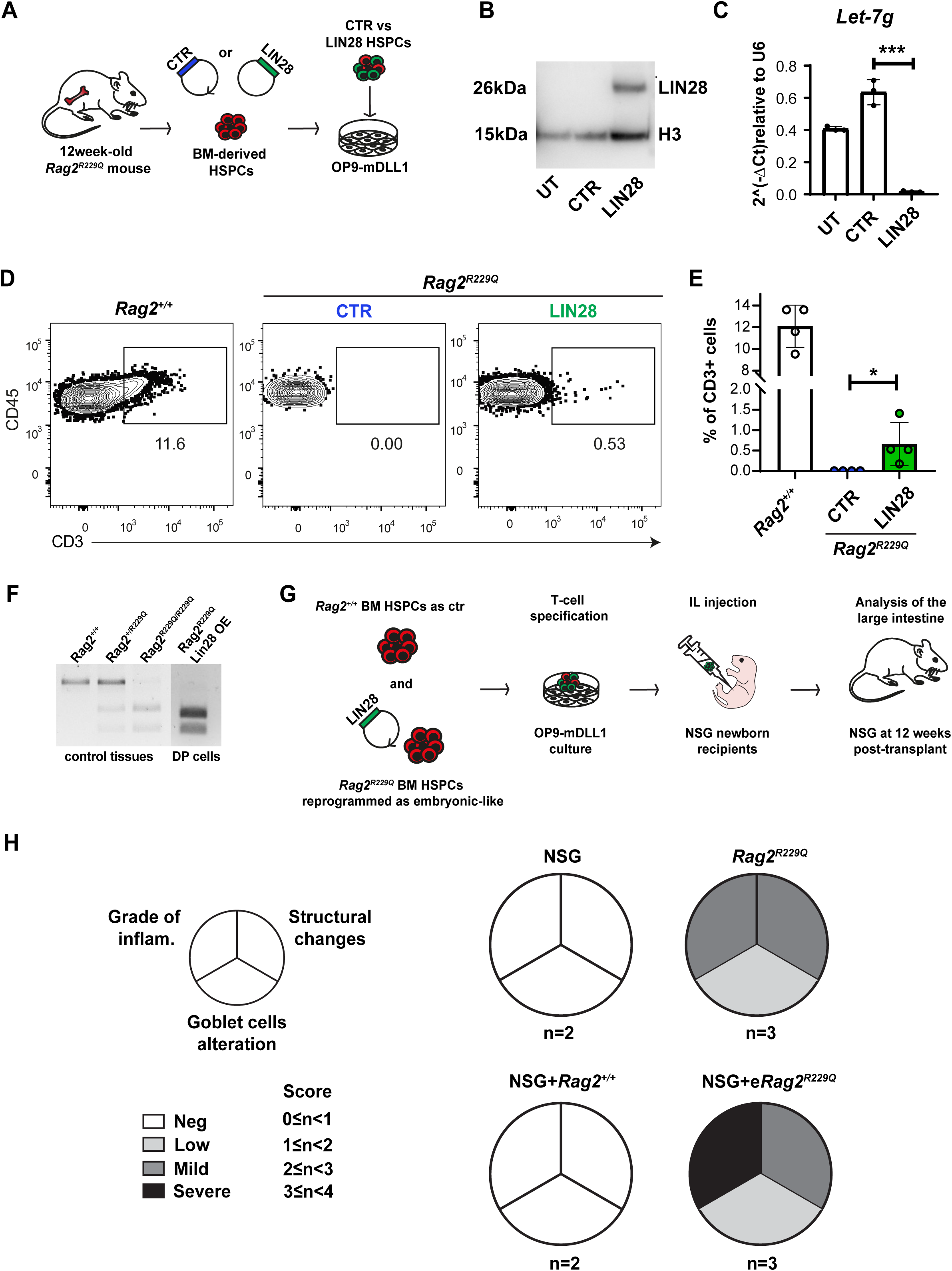
Lin28-reprogrammed embryonic-like *Rag2^R229Q^* HSPCs generate T-cells *ex-vivo* and trigger tissue inflammation in the large intestine of immunodeficient mice. (A) Experimental layout: HSPCs were isolated from the BM of *Rag2^R229Q^* mice, transduced overnight with either a control (CTR) or a LIN28 lentiviral vector, and cultured on OP9-DLL1 stroma for T-cell differentiation. (B) Analysis of LIN28 protein (26kDa) expression by Western Blot in BM-derived HSPCs that were untransduced (UT) or transduced with either a control (CTR) or a LIN28-NGFR lentiviral vector. Histone3 (H3, 15kDa) antibody was chosen as housekeeping control. n=3, independent. (C) qPCR analysis was performed on untransduced (UT), control- and LIN28-transduced CD45^+^ cells to assess the expression of the *Let-7g* miRNA, a known LIN28-target. p-values are for unpaired Student’s t-test. n=3, independent; ***p<0.001. (D) Representative flow cytometric analysis of CD3^+^ T-cells generated from *Rag2^+/+^* (left) and *Rag2^R229Q^* HSPCs transduced with either the control (CTR, middle) or the LIN28 lentiviral vector (right). Cells were gated on SSC/FSC/7AAD^neg^/CD45^+^ cells. (E) Quantification of CD3^+^ T-cells present in *Rag2^+/+^*(white) and *Rag2^R229Q^* HSPCs transduced with either the control (blue) or the LIN28-NGFR lentiviral vector (green). (F) Agarose gel showing the PCR genotyping analysis of LIN28-rescued CD4^+^CD8^+^ T-cells isolated from end-stage cultures. Control DNA from *Rag2^+/+^*, heterozygous and homozygous samples are also shown for comparison. (G) Experimental layout: T-cell progenitors generated from embryonic-like *Rag2^R229Q^* progenitors upon OP9-DLL1 coculture were FAC-sorted for CD45 and injected into the liver on NSG newborns (post-natal day 3). The same mice were analyzed 12 weeks post-transplant to assess the presence of CD3^+^ infiltrates and peripheral inflammation. (H) Pie charts showing the inflammatory score of NSG, *Rag2^R229Q^* control mice, and NSG mice transplanted with *Rag2^+/+^*and *Rag2^R229Q^* embryonic-like T-cell progenitors, determined as explained in the Methods section. Negative: 0<Inf.Score>1; Low: 1<Inf.Score>2; Milld: 2<Inf.Score>3; Severe: 3<Inf.Score>4.

**Supplementary Figure 4.**
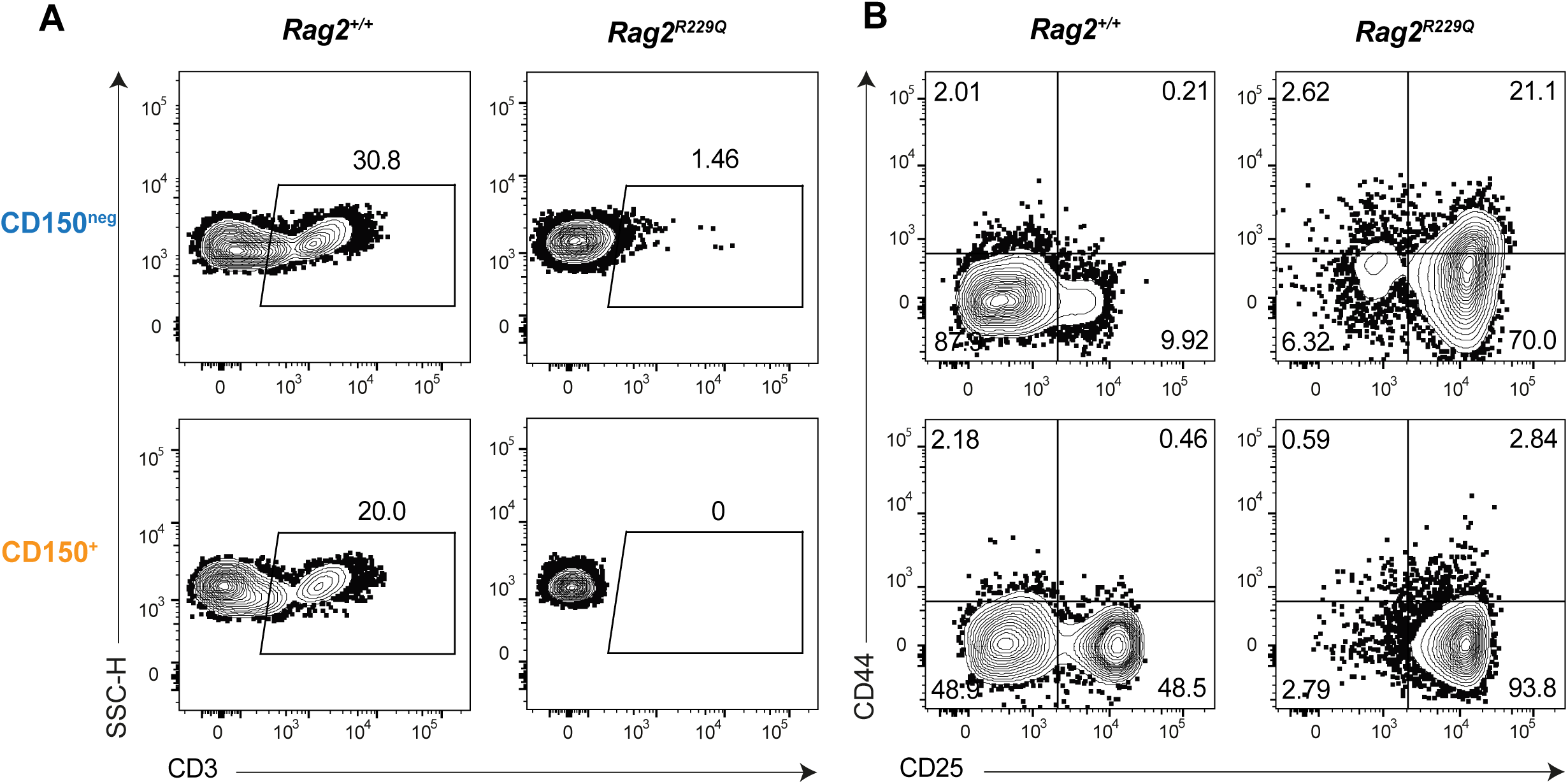
*Rag2^R229Q^*T-cell potential is restricted to HSC-independent embryonic progenitors. (A) Representative flow cytometric analysis of CD3^+^ T-cells output from CD150^neg^ or CD150^pos^ cells isolated from SSC/FSC/7AADneg/Lin^neg^/cKIT^+^/SCA-1^+^population. (B) The progression of E14.5 FL CD150^neg^ or CD150^pos^ progenitors from double negative (DN) stage 1 to 4 was assessed by flow cytometric analysis of CD25 and CD44 expression. Representative plots are shown. Gating strategy shown in Figure S5F.

**Supplementary Figure 5.**
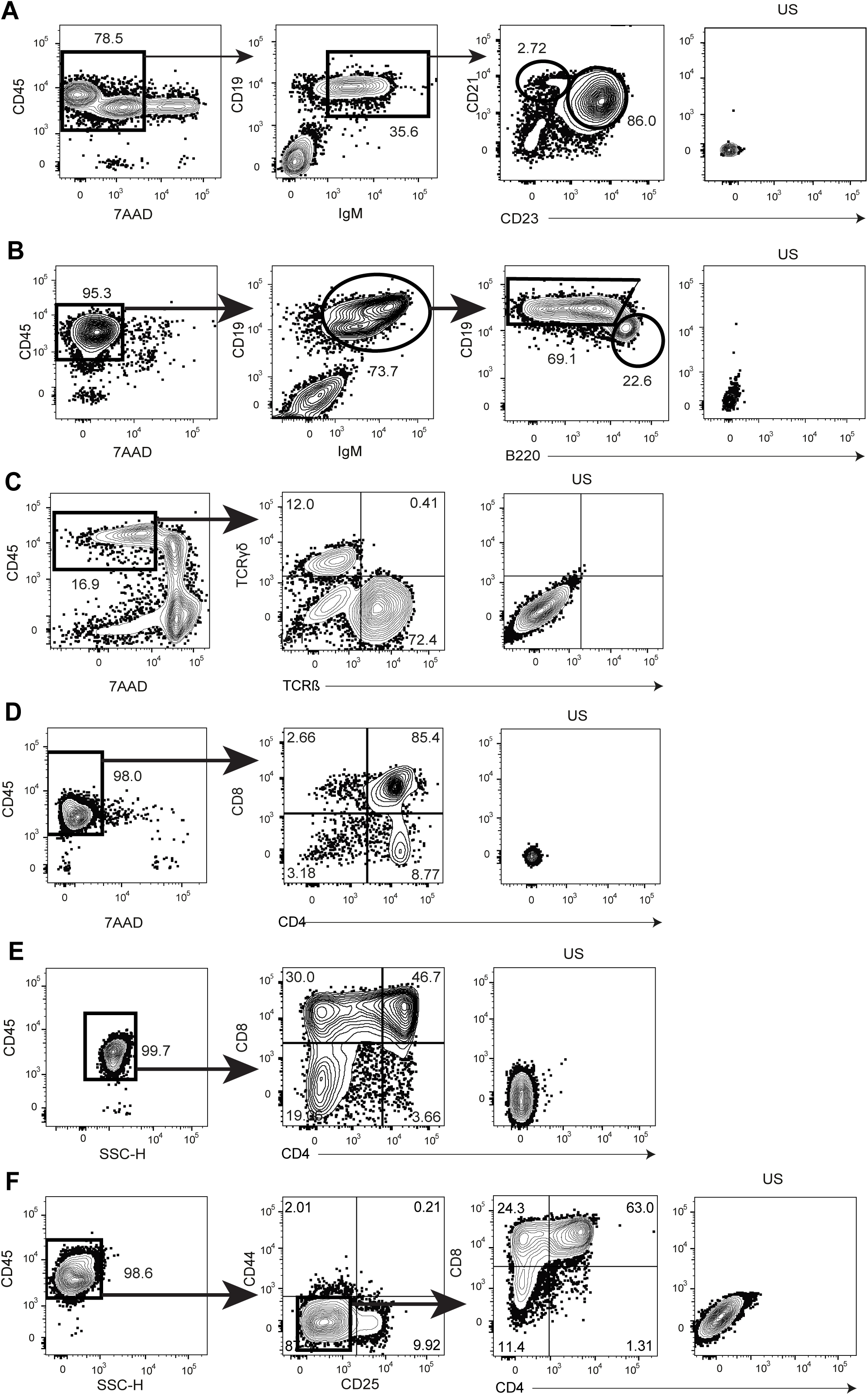
Gating strategies for FACS analysis. (A) Analysis of MZ and FO B-cells in the spleen. First panel: gated on SSC/FSC. Second panel: gated on SSC/FSC/CD45^+^/7AAD^neg^. Third panel: gated on SSC/FSC/Live/CD45^+^/CD19^+^/IgM^+^. Fourth panel: unstained control (US). (B) Analysis of B1/B2 B-cells in the peritoneal cavities. First panel: gated on SSC/FSC. Second panel: gated on SSC/FSC/CD45^+^/7AAD^neg^. Third panel: gated on SSC/FSC/Live/CD45^+^/CD19^+^/IgM^+^. Fourth panel: unstained control (US). (C) Analysis of peripheral T-cells in the large intestine. First panel: gated on SSC/FSC. Second panel: gated on SSC/FSC/CD45^+^/7AAD^neg^. Third panel: unstained control (US). (D) Analysis of murine thymocytes. First panel: gated on SSC/FSC. Second panel: gated on SSC/FSC/CD45^+^/7AAD^neg^. Third panel: unstained control (US). (E and F) Representative flow cytometric analysis showing the gating strategy to analyze murine T-cell potential *in vitro*. (E) Adult T-cells. First panel: gated on SSC/FSC. Second panel: gated on SSC/FSC/CD45^+^. Third panel: unstained control (US). (F) Embryonic T-cells. First panel: gated on SSC/FSC. Second panel: gated on SSC/FSC/CD45^+^. Third panel: SSC/FSC/CD45^+^/CD44^neg^/CD25^neg^. Fourth panel: unstained control (US).

## References

1. Fischer, A., Notarangelo, L.D., Neven, B., Cavazzana, M., and Puck, J.M. (2015). Severe combined immunodeficiencies and related disorders. Nat Rev Dis Primers 1, 15061. 10.1038/nrdp.2015.61.

2. Villa, A., Notarangelo, L.D., and Roifman, C.M. (2008). Omenn syndrome: inflammation in leaky severe combined immunodeficiency. J Allergy Clin Immunol 122, 1082–1086. 10.1016/j.jaci.2008.09.037.

3. Sharapova, S.O., Guryanova, I.E., Pashchenko, O.E., Kondratenko, I.V., Kostyuchenko, L.V., Rodina, Y.A., Varlamova, T.V., Bondarenko, A.V., Chernyshova, L.I., Gyseva, M.N., et al. (2015). Molecular Characteristics, Clinical and Immunologic Manifestations of 11 Children with Omenn Syndrome in East Slavs (Russia, Belarus, Ukraine). J Clin Immunol 36, 46–55. 10.1007/s10875-015-0216-7.

4. Gruber, T.A., Shah, A.J., Hernandez, M., Crooks, G.M., Abdel-Azim, H., Gupta, S., McKnight, S., White, D., Kapoor, N., and Kohn, D.B. (2009). Clinical and genetic heterogeneity in Omenn syndrome and severe combined immune deficiency. Pediatr Transplant 13, 244–250. 10.1111/j.1399-3046.2008.00970.x.

5. Delmonte, O.M., Villa, A., and Notarangelo, L.D. (2020). Immune dysregulation in patients with RAG deficiency and other forms of combined immune deficiency. Blood 135, 610–619. 10.1182/blood.2019000923.

6. Giliani, S., Bonfim, C., Basile, G. de S., Lanzi, G., Brousse, N., Koliski, A., Malvezzi, M., Fischer, A., Notarangelo, L.D., and Deist, F.L. (2006). Omenn syndrome in an infant with IL7RA gene mutation. J Pediatrics 148, 272–274. 10.1016/j.jpeds.2005.10.004.

7. Schuetz, C., Gerke, J., Ege, M., Walter, J., Kusters, M., Worth, A., Kanakry, J.A., Dimitrova, D., Wolska-Kuśnierz, B., Chen, K., et al. (2023). Hypomorphic RAG deficiency: impact of disease burden on survival and thymic recovery argues for early diagnosis and HSCT. Blood 141, 713–724. 10.1182/blood.2022017667.

8. Rigoni, R., Fontana, E., Dobbs, K., Marrella, V., Taverniti, V., Maina, V., Facoetti, A., D’Amico, G., Al-Herz, W., Cruz-Munoz, M.E., et al. (2020). Cutaneous barrier leakage and gut inflammation drive skin disease in Omenn syndrome. J Allergy Clin Immun 146, 1165–1179.e11. 10.1016/j.jaci.2020.04.005.

9. Rieux-Laucat, F., Bahadoran, P., Brousse, N., Selz, F., Fischer, A., Deist, F.L., and Villartay, J.P.D. (1998). Highly restricted human T cell repertoire in peripheral blood and tissue-infiltrating lymphocytes in Omenn’s syndrome. J Clin Invest 102, 312–321. 10.1172/jci332.

10. Zhang, J., Quintal, L., Atkinson, A., Williams, B., Grunebaum, E., and Roifman, C.M. (2005). Novel RAG1 Mutation in a Case of Severe Combined Immunodeficiency. Pediatrics 116, e445–e449. 10.1542/peds.2005-0369.

11. Signorini, S., Imberti, L., Pirovano, S., Villa, A., Facchetti, F., Ungari, M., Bozzi, F., Albertini, A., Ugazio, A.G., Vezzoni, P., et al. (1999). Intrathymic Restriction and Peripheral Expansion of the T-Cell Repertoire in Omenn Syndrome. Blood 94, 3468–3478. 10.1182/blood.v94.10.3468.422k34_3468_3478.

12. Lev, A., Simon, A.J., Amariglio, N., Rechavi, G., and Somech, R. (2012). Selective clinical and immune response of the oligoclonal autoreactive T cells in Omenn patients after cyclosporin A treatment. Clin Exp Immunol 167, 338–345. 10.1111/j.1365-2249.2011.04508.x.

13. Min, Q., Csomos, K., Li, Y., Dong, L., Hu, Z., Meng, X., Yu, M., Walter, J.E., and Wang, J.-Y. (2023). B cell abnormalities and autoantibody production in patients with partial RAG deficiency. Front. Immunol. 14, 1155380. 10.3389/fimmu.2023.1155380.

14. Ghosn, E., Yoshimoto, M., Nakauchi, H., Weissman, I.L., and Herzenberg, L.A. (2019). Hematopoietic stem cell-independent hematopoiesis and the origins of innate-like B lymphocytes. Development 146, dev170571. 10.1242/dev.170571.

15. Bendelac, A., Bonneville, M., and Kearney, J.F. (2001). Autoreactivity by design: innate B and T lymphocytes. Nat Rev Immunol 1, 177–186. 10.1038/35105052.

16. Suo, C., Dann, E., Goh, I., Jardine, L., Kleshchevnikov, V., Park, J.-E., Botting, R.A., Stephenson, E., Engelbert, J., Tuong, Z.K., et al. (2022). Mapping the developing human immune system across organs. Biorxiv, 2022.01.17.476665. 10.1101/2022.01.17.476665.

17. Papadopoulou, M., Tieppo, P., McGovern, N., Gosselin, F., Chan, J.K.Y., Goetgeluk, G., Dauby, N., Cogan, A., Donner, C., Ginhoux, F., et al. (2019). TCR Sequencing Reveals the Distinct Development of Fetal and Adult Human Vγ9Vδ2 T Cells. J Immunol 203, 1468–1479. 10.4049/jimmunol.1900592.

18. Carvalho, T.L., Mota-Santos, T., Cumano, A., Demengeot, J., and Vieira, P. (2001). Arrested B Lymphopoiesis and Persistence of Activated B Cells in Adult Interleukin 7−/− Mice. J Exp Med 194, 1141–1150. 10.1084/jem.194.8.1141.

19. Mass, E., Jacome-Galarza, C.E., Blank, T., Lazarov, T., Durham, B.H., Ozkaya, N., Pastore, A., Schwabenland, M., Chung, Y.R., Rosenblum, M.K., et al. (2017). A somatic mutation in erythro-myeloid progenitors causes neurodegenerative disease. Nature 549, 389–393. 10.1038/nature23672.

20. Symeonidou, V., Jakobczyk, H., Bashanfer, S., Malouf, C., Fotopoulou, F., Kotecha, R.S., Anderson, R.A., Finch, A.J., and Ottersbach, K. (2021). Defining the fetal origin of MLL-AF4 infant leukemia highlights specific fatty acid requirements. Cell Reports 37, 109900. 10.1016/j.celrep.2021.109900.

21. Yoshimoto, M., Montecino-Rodriguez, E., Ferkowicz, M.J., Porayette, P., Shelley, W.C., Conway, S.J., Dorshkind, K., and Yoder, M.C. (2011). Embryonic day 9 yolk sac and intra-embryonic hemogenic endothelium independently generate a B-1 and marginal zone progenitor lacking B-2 potential. P Natl Acad Sci Usa 108, 1468–1473. 10.1073/pnas.1015841108.

22. Kikuchi, K., and Kondo, M. (2006). Developmental switch of mouse hematopoietic stem cells from fetal to adult type occurs in bone marrow after birth. Proc National Acad Sci 103, 17852–17857. 10.1073/pnas.0603368103.

23. Kobayashi, M., Shelley, W.C., Seo, W., Vemula, S., Lin, Y., Liu, Y., Kapur, R., Taniuchi, I., and Yoshimoto, M. (2014). Functional B-1 progenitor cells are present in the hematopoietic stem cell-deficient embryo and depend on Cbfβ for their development. P Natl Acad Sci Usa 111, 12151–12156. 10.1073/pnas.1407370111.

24. Gentek, R., Ghigo, C., Hoeffel, G., Jorquera, A., Msallam, R., Wienert, S., Klauschen, F., Ginhoux, F., and Bajénoff, M. (2018). Epidermal γδ T cells originate from yolk sac hematopoiesis and clonally self-renew in the adult. J Exp Medicine 215, 2994–3005. 10.1084/jem.20181206.

25. Sandrock, I., Reinhardt, A., Ravens, S., Binz, C., Wilharm, A., Martins, J., Oberdörfer, L., Tan, L., Lienenklaus, S., Zhang, B., et al. (2018). Genetic models reveal origin, persistence and non-redundant functions of IL-17–producing γδ T cells. J Exp Med 215, 3006–3018. 10.1084/jem.20181439.

26. Haas, J.D., Ravens, S., Düber, S., Sandrock, I., Oberdörfer, L., Kashani, E., Chennupati, V., Föhse, L., Naumann, R., Weiss, S., et al. (2012). Development of Interleukin-17-Producing γδ T Cells Is Restricted to a Functional Embryonic Wave. Immunity 37, 48–59. 10.1016/j.immuni.2012.06.003.

27. Lee, Y.T., Vasconcellos, J.F. de, Yuan, J., Byrnes, C., Noh, S.-J., Meier, E.R., Kim, K.S., Rabel, A., Kaushal, M., Muljo, S.A., et al. (2013). LIN28B-mediated expression of fetal hemoglobin and production of fetal-like erythrocytes from adult human erythroblasts ex vivo. Blood 122, 1034–1041. 10.1182/blood-2012-12-472308.

28. Stolla, M.C., Catherman, S.C., Kingsley, P.D., Rowe, R.G., Koniski, A.D., Fegan, K., Vit, L., McGrath, K.E., Daley, G.Q., and Palis, J. (2019). Lin28b regulates age-dependent differences in murine platelet function. Blood Adv 3, 72–82. 10.1182/bloodadvances.2018020859.

29. Yuan, J., Nguyen, C.K., Liu, X., Kanellopoulou, C., and Muljo, S.A. (2012). Lin28b reprograms adult bone marrow hematopoietic progenitors to mediate fetal-like lymphopoiesis. Sci New York N Y 335, 1195–1200. 10.1126/science.1216557.

30. Marrella, V., Poliani, P.L., Casati, A., Rucci, F., Frascoli, L., Gougeon, M.-L., Lemercier, B., Bosticardo, M., Ravanini, M., Battaglia, M., et al. (2007). A hypomorphic R229Q Rag2 mouse mutant recapitulates human Omenn syndrome. J Clin Invest 117, 1260–1269. 10.1172/jci30928.

31. Cassani, B., Poliani, P.L., Marrella, V., Schena, F., Sauer, A.V., Ravanini, M., Strina, D., Busse, C.E., Regenass, S., Wardemann, H., et al. (2010). Homeostatic expansion of autoreactive immunoglobulin-secreting cells in the Rag2 mouse model of Omenn syndrome. J Exp Medicine 207, 1525–1540. 10.1084/jem.20091928.

32. Ghosn, E.E.B., Yamamoto, R., Hamanaka, S., Yang, Y., Herzenberg, L.A., Nakauchi, H., and Herzenberg, L.A. (2012). Distinct B-cell lineage commitment distinguishes adult bone marrow hematopoietic stem cells. Proc. Natl. Acad. Sci. 109, 5394–5398. 10.1073/pnas.1121632109.

33. Harville, T.O., Adams, D.M., Howard, T.A., and Ware, R.E. (1997). Oligoclonal Expansion of CD45RO+ T Lymphocytes in Omenn Syndrome. J Clin Immunol 17, 322–332. 10.1023/a:1027330800085.

34. Villa, A. (2011). Omenn Syndrome: inflammation and autoimmunity. J Transl Med 9, I5. 10.1186/1479-5876-9-s2-i5.

35. Lee, Y.N., Frugoni, F., Dobbs, K., Tirosh, I., Du, L., Ververs, F.A., Ru, H., Bruin, L.O. de, Adeli, M., Bleesing, J.H., et al. (2016). Characterization of T and B cell repertoire diversity in patients with RAG deficiency. Sci. Immunol. 1. 10.1126/sciimmunol.aah6109.

36. Holmes, R., and Zúñiga-Pflücker, J.C. (2009). The OP9-DL1 System: Generation of T-Lymphocytes from Embryonic or Hematopoietic Stem Cells In Vitro. Cold Spring Harb Protoc 2009, pdb.prot5156. 10.1101/pdb.prot5156.

37. Yoshimoto, M., Porayette, P., Glosson, N.L., Conway, S.J., Carlesso, N., Cardoso, A.A., Kaplan, M.H., and Yoder, M.C. (2012). Autonomous murine T-cell progenitor production in the extra-embryonic yolk sac before HSC emergence. Blood 119, 5706–5714. 10.1182/blood-2011-12-397489.

38. Viswanathan, S.R., Daley, G.Q., and Gregory, R.I. (2008). Selective Blockade of MicroRNA Processing by Lin28. Science 320, 97–100. 10.1126/science.1154040.

39. Kolar, G.R., Yokota, T., Rossi, M.I.D., Nath, S.K., and Capra, J.D. (2004). Human fetal, cord blood, and adult lymphocyte progenitors have similar potential for generating B cells with a diverse immunoglobulin repertoire. Blood 104, 2981–2987. 10.1182/blood-2003-11-3961.

40. Adkins, B. (2003). Peripheral CD4+ Lymphocytes Derived from Fetal versus Adult Thymic Precursors Differ Phenotypically and Functionally. J. Immunol. 171, 5157–5164. 10.4049/jimmunol.171.10.5157.

41. Igarashi, H., Kouro, T., Yokota, T., Comp, P.C., and Kincade, P.W. (2001). Age and stage dependency of estrogen receptor expression by lymphocyte precursors. Proc. Natl. Acad. Sci. 98, 15131–15136. 10.1073/pnas.011513098.

42. Rackaityte, E., and Halkias, J. (2020). Mechanisms of Fetal T Cell Tolerance and Immune Regulation. Front. Immunol. 11, 588. 10.3389/fimmu.2020.00588.

43. Li, Y., Innocentin, S., Withers, D.R., Roberts, N.A., Gallagher, A.R., Grigorieva, E.F., Wilhelm, C., and Veldhoen, M. (2011). Exogenous Stimuli Maintain Intraepithelial Lymphocytes via Aryl Hydrocarbon Receptor Activation. Cell 147, 629–640. 10.1016/j.cell.2011.09.025.

44. Cumano, A., Ferraz, J.C., Klaine, M., Santo, J.P.D., and Godin, I. (2001). Intraembryonic, but Not Yolk Sac Hematopoietic Precursors, Isolated before Circulation, Provide Long-Term Multilineage Reconstitution. Immunity 15, 477–485. 10.1016/s1074-7613(01)00190-x.

45. Ghosn, E.E.B., Waters, J., Phillips, M., Yamamoto, R., Long, B.R., Yang, Y., Gerstein, R., Stoddart, C.A., Nakauchi, H., and Herzenberg, L.A. (2015). Fetal Hematopoietic Stem Cell Transplantation Fails to Fully Regenerate the B-Lymphocyte Compartment. Stem Cell Rep 6, 137–149. 10.1016/j.stemcr.2015.11.011.

46. Parekh, C., and Crooks, G.M. (2013). Critical Differences in Hematopoiesis and Lymphoid Development between Humans and Mice. J Clin Immunol 33, 711–715. 10.1007/s10875-012-9844-3.

47. Dasouki, M., Jabr, A., AlDakheel, G., Elbadaoui, F., Alazami, A.M., Al-Saud, B., Arnaout, R., Aldhekri, H., Alotaibi, I., Al-Mousa, H., et al. (2020). TREC and KREC profiling as a representative of thymus and bone marrow output in patients with various inborn errors of immunity. Clin Exp Immunol 202, 60–71. 10.1111/cei.13484.

48. Sturgeon, C.M., Ditadi, A., Awong, G., Kennedy, M., and Keller, G. (2014). Wnt Signaling Controls the Specification of Definitive and Primitive Hematopoiesis From Human Pluripotent Stem Cells. Nat. Biotechnol. 32, 554–561. 10.1038/nbt.2915.

49. Ditadi, A., and Sturgeon, C.M. (2016). Directed differentiation of definitive hemogenic endothelium and hematopoietic progenitors from human pluripotent stem cells. Methods 101, 65–72. 10.1016/j.ymeth.2015.10.001.

50. Ditadi, A., Sturgeon, C.M., Tober, J., Awong, G., Kennedy, M., Yzaguirre, A.D., Azzola, L., Ng, E.S., Stanley, E.G., French, D.L., et al. (2015). Human definitive haemogenic endothelium and arterial vascular endothelium represent distinct lineages. Nat Cell Biol 17, 580–591. 10.1038/ncb3161.

51. Luff, S.A., Creamer, J.P., Valsoni, S., Dege, C., Scarfò, R., Dacunto, A., Cascione, S., Randolph, L.N., Cavalca, E., Merelli, I., et al. (2022). Identification of a retinoic acid-dependent haemogenic endothelial progenitor from human pluripotent stem cells. Nat Cell Biol, 1–9. 10.1038/s41556-022-00898-9.

52. Montel-Hagen, A., Seet, C.S., Li, S., Chick, B., Zhu, Y., Chang, P., Tsai, S., Sun, V., Lopez, S., Chen, H.-C., et al. (2019). Organoid-Induced Differentiation of Conventional T Cells from Human Pluripotent Stem Cells. Cell Stem Cell 24, 376–389.e8. 10.1016/j.stem.2018.12.011.

53. Bosticardo, M., Pala, F., Calzoni, E., Delmonte, O.M., Dobbs, K., Gardner, C.L., Sacchetti, N., Kawai, T., Garabedian, E.K., Draper, D., et al. (2020). Artificial thymic organoids represent a reliable tool to study T-cell differentiation in patients with severe T-cell lymphopenia. Blood Adv. 4, 2611– 2616. 10.1182/bloodadvances.2020001730.

54. Brauer, P.M., Pessach, I.M., Clarke, E., Rowe, J.H., Bruin, L.O. de, Lee, Y.N., Dominguez-Brauer, C., Comeau, A.M., Awong, G., Felgentreff, K., et al. (2016). Modeling altered T-cell development with induced pluripotent stem cells from patients with RAG1-dependent immune deficiencies. Blood 128, 783–793. 10.1182/blood-2015-10-676304.

55. Bifsha, P., Leiding, J.W., Pai, S.-Y., Colamartino, A.B.L., Hartog, N., Church, J.A., Oshrine, B.R., Puck, J.M., Markert, M.L., and Haddad, E. (2020). Diagnostic assay to assist clinical decisions for unclassified severe combined immune deficiency. Blood Adv. 4, 2606–2610. 10.1182/bloodadvances.2020001736.

56. Cassani, B., Poliani, P.L., Moratto, D., Sobacchi, C., Marrella, V., Imperatori, L., Vairo, D., Plebani, A., Giliani, S., Vezzoni, P., et al. (2010). Defect of regulatory T cells in patients with Omenn syndrome. J Allergy Clin Immunol 125, 209–216. 10.1016/j.jaci.2009.10.023.

57. Cavadini, P., Vermi, W., Facchetti, F., Fontana, S., Nagafuchi, S., Mazzolari, E., Sediva, A., Marrella, V., Villa, A., Fischer, A., et al. (2005). AIRE deficiency in thymus of 2 patients with Omenn syndrome. J Clin Invest 115, 728–732. 10.1172/jci23087.

58. Somech, R., Simon, A.J., Lev, A., Dalal, I., Spirer, Z., Goldstein, I., Nagar, M., Amariglio, N., Rechavi, G., and Roifman, C.M. (2009). Reduced central tolerance in Omenn syndrome leads to immature self-reactive oligoclonal T cells. J Allergy Clin Immun 124, 793–800. 10.1016/j.jaci.2009.06.048.

59. Rother, M.B., Jensen, K., Burg, M. van der, Bovenkamp, F.S. van de, Kroek, R., IJcken, W.F.J. van, Velden, V.H.J. van der, Cupedo, T., Olstad, O.K., Dongen, J.J.M. van, et al. (2016). Decreased IL7Rα and TdT expression underlie the skewed immunoglobulin repertoire of human B-cell precursors from fetal origin. Sci Rep-uk 6, 33924. 10.1038/srep33924.

60. The regulated expression of B lineage associated genes during B cell differentiation in bone marrow and fetal liver (1993). J Exp Medicine 178, 951–960. 10.1084/jem.178.3.951.

61. Rechavi, E., Lev, A., Lee, Y.N., Simon, A.J., Yinon, Y., Lipitz, S., Amariglio, N., Weisz, B., Notarangelo, L.D., and Somech, R. (2015). Timely and spatially regulated maturation of B and T cell repertoire during human fetal development. Sci Transl Med 7, 276ra25–276ra25. 10.1126/scitranslmed.aaa0072.

62. Ponchel, F., Morgan, A.W., Bingham, S.J., Quinn, M., Buch, M., Verburg, R.J., Henwood, J., Douglas, S.H., Masurel, A., Conaghan, P., et al. (2002). Dysregulated lymphocyte proliferation and differentiation in patients with rheumatoid arthritis. Blood 100, 4550–4556. 10.1182/blood-2002-03-0671.

63. Waase, I., Kayser, C., Carlson, P.J., Goronzy, J.J., and Weyand, C.M. (1996). Oligoclonal T cell proliferation in patients with rheumatoid arthritis and their unaffected siblings. Arthritis Rheumatism 39, 904–913. 10.1002/art.1780390606.

64. Gomez-Tourino, I., Kamra, Y., Baptista, R., Lorenc, A., and Peakman, M. (2017). T cell receptor β-chains display abnormal shortening and repertoire sharing in type 1 diabetes. Nat Commun 8, 1792. 10.1038/s41467-017-01925-2.

65. Sakkas, L.I., Xu, B., Artlett, C.M., Lu, S., Jimenez, S.A., and Platsoucas, C.D. (2002). Oligoclonal T Cell Expansion in the Skin of Patients with Systemic Sclerosis. J Immunol 168, 3649–3659. 10.4049/jimmunol.168.7.3649.

66. Dolens, A.-C., Durinck, K., Lavaert, M., Meulen, J.V. der, Velghe, I., Medts, J.D., Weening, K., Roels, J., Mulder, K.D., Volders, P.-J., et al. (2020). Distinct Notch1 and BCL11B requirements mediate human γδ/αβ T cell development. Embo Rep 21, e49006. 10.15252/embr.201949006.

